# Sustained loss of *Pptc7* triggers variable skeletal muscle dysfunction and diminished body mass through dysregulation of BNIP3

**DOI:** 10.64898/2026.07.13.738264

**Authors:** Tessa M. Lochetto, Thiago N. Menezes, Kevin Cho, Srikar Vegesna, Michaela M. Morhaus, Jeremie L. A. Ferey, Merima Forny, Leah P. Shriver, Gretchen A. Meyer, Gary J. Patti, Natalie M. Niemi

## Abstract

The mitochondrial phosphatase PPTC7 is required to sustain mammalian metabolism, as its global knockout (KO) triggers hypoketotic hypoglycemia and perinatal lethality in mice. However, the extent to which the loss of *Pptc7* manifests pathology beyond the perinatal transition is unknown. Furthermore, PPTC7 was recently identified as dual functional, regulating mitochondrial protein phosphorylation and receptor mediated mitophagy, rendering it unclear which function(s) may influence in vivo physiology. Here, we find that sustained, inducible *Pptc7* KO decreased lean mass, compromised whole body oxygen consumption, and altered circulating metabolites in adult male mice. We hypothesized that these phenotypes stemmed from skeletal muscle dysfunction and found lower mass and fiber cross-sectional area with shifts in fiber type distribution in select muscles of the hindlimb in *Pptc7* KO animals. Loss of PPTC7 increased BNIP3 protein levels and decreased mitochondrial content in skeletal muscle, suggesting elevated mitophagy may drive pathology. Consistently, KO of *Bnip3* rescued the lower body weight and lean mass seen in inducible *Pptc7* KO adult animals and partially rescued perinatal lethality in global *Pptc7* KO mice. These data demonstrate that loss of PPTC7 incites surprisingly variable dysfunction across physiological contexts that at least partially stems from dysregulated BNIP3.

## Introduction

Mitochondria are essential organelles present in virtually all eukaryotic cells. Beyond their canonical roles as the “powerhouses of the cell,” mitochondria act as cellular metabolic hubs through the coordination of fatty acid oxidation, amino acid catabolism, redox biochemistry, heme biosynthesis, and calcium homeostasis, amongst other pathways (*1*, *2*). Though well-established to perform these diverse roles, the mechanisms underlying the systems-level regulation of these mitochondrial functions are not well defined, particularly across tissue types and in response to diverse cellular and organismal demands.

One emerging paradigm for mitochondrial metabolic regulation is dynamic protein phosphorylation. Recent evidence suggests that most proteins within mitochondria can be phosphorylated (*3*), yet the extent to which most of these post-translational modifications regulate mitochondrial pathways–or even have functional significance–remains unclear. One piece of evidence suggesting that phosphorylation may be broadly meaningful for mitochondrial function is the presence of multiple phosphatases at or within these organelles in organisms ranging from yeast to mammals (*3–7*). These phosphatases reside across mitochondrial compartments (*3*) and their presence suggests that, minimally, dephosphorylation of select molecules enables organellar homeostasis. Previously, we demonstrated that genetic ablation of the mitochondrial phosphatase PPTC7, or its yeast ortholog Ptc7p, reproducibly increased the occupancy of phosphorylation sites on multiple mitochondrial proteins, consistent with its function as a protein phosphatase (*4–7*). Strikingly, mice with genetically ablated *Pptc7* displayed robust metabolic dysfunction, including hypoglycemic hypoketosis with lactic acid accumulation that culminated in fully penetrant lethality within one day of birth (*6*). These initial data pointed to a critical yet underappreciated role of mitochondrial protein phosphorylation in orchestrating metabolic regulation in mice, yet the specific molecular drivers of the perinatal lethal phenotype were not fully defined.

However, the interpretation that dysregulated protein phosphorylation may drive pathophysiology in *Pptc7* KO systems has been complicated by the fact that PPTC7 was recently discovered to have a second role as a negative regulator of BNIP3- and NIX-mediated mitophagy (*7–10*). While PPTC7 and Ptc7p have been mapped to the mitochondrial matrix using unbiased proteomics approaches (*11*, *12*), we and others have identified a second population of PPTC7 that localizes to the outer mitochondrial membrane (OMM) (*8–10*). Here, PPTC7 bridges the mitophagy receptors BNIP3 and NIX to the E3 ubiquitin ligase FBXL4 to facilitate their proteasomal turnover (*9*, *10*, *13–16*). As such, loss of either FBXL4 or PPTC7 results in unchecked and excessive levels of BNIP3/NIX-mediated mitophagy, leading to a robust loss of mitochondrial content across cell types and tissues. Importantly, ablation of BNIP3 or NIX expression in *Fbxl4* or *Pptc7* knockout (KO) systems reverses this elevated mitophagic flux and largely rescued mitochondrial content (*7*, *13–15*), demonstrating that FBXL4 and PPTC7 specifically regulate the receptor-mediated mitophagic pathway.

Given their proximal roles in the molecular regulation of BNIP3 and NIX, it is reasonable to assume that loss of either PPTC7 or FBXL4 would trigger similar physiological phenotypes. Indeed, inactivating mutations in *FBXL4* or *PPTC7* elicit severe, typically early-onset mitochondrial disease in humans, characterized by metabolic acidosis, pronounced hypotonia, neurological dysfunction, and broad developmental delays (*17–19*). Furthermore, a global knockout *Fbxl4* mouse model shares multiple phenotypes with *Pptc7* KO animals, including substantial, though not fully penetrant, perinatal lethality (*20*). This lethality could be rescued by ablation of either *Bnip3* or *Bnip3l* (gene name for NIX) within the *Fbxl4* KO background (*13*), and analysis of liver tissue from perinatal *Fbxl4* KO animals revealed similar metabolic derangements as seen in perinatal *Pptc7* KO animals (*6*, *13*). Given the substantial overlap in phenotypes amongst these mouse models, these data suggest that much of the pathology manifested in the perinatal period upon loss of *Pptc7* likely derives from the dysregulation of BNIP3- and NIX-mediated mitophagy. However, the extent to which loss of PPTC7 induces pathology beyond the perinatal transition in animal models is relatively unexplored. Furthermore, given the dual functional role of PPTC7 in mitochondrial biology, establishing whether pathologies arising due to loss of *Pptc7* stem from dysregulated protein phosphorylation (i.e., intrinsic organellar dysfunction), decreased mitochondrial content (i.e., excessive and unchecked mitophagy), or a combination of these pathways will be critical to understand how this mitochondrial phosphatase regulates mammalian metabolism and physiology.

Here, we exploit our recently generated conditional *Pptc7* mouse model (*7*) to characterize the physiological and pathophysiological effects of chronic loss of *Pptc7* in adult mice. Though our previous global *Pptc7* KO model displayed fully penetrant perinatal lethality, we found that induction of sustained *Pptc7* KO in adult animals is compatible with viability. However, we find various physiological abnormalities in *Pptc7* KO animals, including lower body mass, metabolic derangements, and selective skeletal muscle dysfunction. As PPTC7 has dual functions, we investigated mitochondria isolated from *Pptc7* KO animals to probe the potential mechanism(s) driving skeletal muscle dysfunction. Importantly, these mitochondria retained respiratory capacity when normalized for content defects, suggesting that a loss of mitochondrial mass drives skeletal muscle dysfunction rather than intrinsic organellar dysregulation. These data suggest that excessive mitophagy induces physiological dysfunction, which we tested by generating *Pptc7*/*Bnip3* double knockout animal models. Consistently, *Bnip3* genetic ablation reversed body weight defects and the loss of lean mass in adult inducible *Pptc7* KO mice and rescued perinatal lethality in germline *Pptc7* KO animals. Together, our data suggest that loss of PPTC7 causes developmentally distinct and tissue-specific dysfunction in mice, which is at least partially caused by diminished mitochondrial content and elevated BNIP3 protein levels in adult mice.

## Results

### Loss of Pptc7 diminishes lean body mass in male and female mice but compromises organismal respiration in a sex-specific manner

To overcome the limitations of studying a perinatal lethal mouse model, we recently generated a conditional mouse model that allows tamoxifen-induced knockout (KO) of *Pptc7* across tissues in adult animals (*7*). We aged control (i.e., *Pptc7*^flox/flox^ mice) and experimental (i.e., UBC-Cre-ER^T2^;*Pptc7*^flox/flox^ mice) littermates to 8 weeks of age, after which we tamoxifen treated for 2 weeks to promote excision of exon 3 within the *Pptc7* gene locus. Upon completion of tamoxifen treatment, we aged these animals and noted that *Pptc7* KO mice of each sex had significantly lower body weight 20 weeks post-tamoxifen (Figures 1C, D), though neither male nor female *Pptc7* KO mice displayed differences in body mass at 8 weeks post-tamoxifen treatment (Figures 1A, B). To understand if this lower body mass stemmed from differences in fat or lean mass, we performed body composition analysis on control and *Pptc7* KO animals at both time points. Neither male nor female mice had significant differences in fat or lean mass 8 weeks after tamoxifen-induced *Pptc7* knockout (Figures 1A, B). However, whole-body quantitative NMR (e.g., EchoMRI) analysis revealed significantly lower lean mass in both sexes of *Pptc7* KO mice at 20-weeks post-tamoxifen treatment independent of significant changes in fat mass (Figures 1C, D). These data suggest that PPTC7 expression is required to maintain murine lean mass over time.

**Figure 1:**
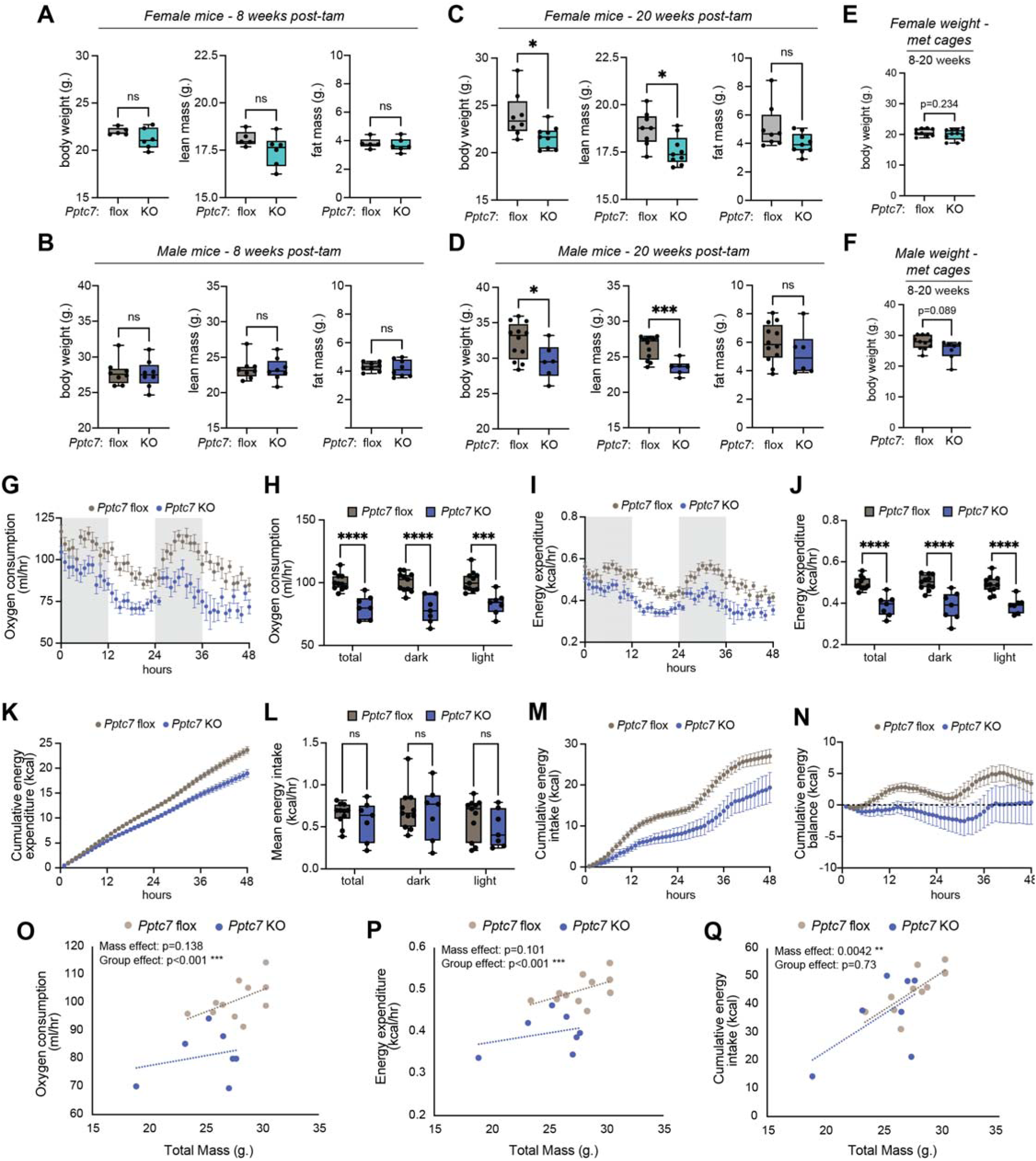
Loss of Pptc7 diminishes lean body mass in male and female mice but compromises organismal respiration in a sex-specific manner. **A.-D**. Total body weight (left), lean mass (center) and fat mass (right) for control (Pptc7 flox) and experimental (Pptc7 knockout, KO) female and male mice at 8 weeks post-tamoxifen treatment (A., B.) and 20-weeks post-tamoxifen treatment (C., D.). n.s. = not significant, *=p<0.05, ***=p<0.001, Welch’s t test. **E**.**F**., Total body weight for female (E.) and male (F.) mice aged 8-20 weeks post-tamoxifen treatment for analysis in metabolic cages. **G**., **H**. Oxygen consumption rates for control (Pptc7 flox) and experimental (Pptc7 KO) male animals. G. 48-hour average of oxygen consumption rates from male Pptc7 flox (n=11) and Pptc7 KO (n=7) animals. Error bars represent S.E.M. H. Oxygen consumption rates shown in G. analyzed by circadian parameters for total day averages as well as those during the dark and light phase. **I**., **J**. Energy expenditure for control (Pptc7 flox) and experimental (Pptc7 KO) male animals. I. Energy expenditure rates from male Pptc7 flox (n=11) and Pptc7 KO (n=7) animals. Error bars represent S.E.M. J. Energy expenditure rates shown in I. analyzed by circadian parameters for total day averages as well as those during the dark and light phase. ****=p<0.0001, Ordinary Two-way ANOVA. **K**. Cumulative energy expenditure for male Pptc7 flox (n=11) and Pptc7 KO (n=7) animals over a 48-hour period; error bars represent S.E.M. **L**. Mean hourly energy intake rates analyzed by circadian parameters for total day averages as well as those during the dark and light phase. n.s. = not significant, Ordinary Two-way ANOVA. **M**. Cumulative energy intake for male Pptc7 flox (n=11) and Pptc7 KO (n=7) animals over a 48-hour period; error bars represent S.E.M. **N**. Cumulative energy balance for male Pptc7 flox (n=11) and Pptc7 KO (n=7) animals over a 48-hour period; error bars represent S.E.M. **O**.-**Q**. ANCOVA analysis of weight-dependent parameters such as oxygen consumption rates (O.), energy expenditure rates (P.), and cumulative energy intake (Q.) for Pptc7 flox (n=11) and Pptc7 KO (n=7) male mice. Statistical analysis performed through the CalR2 app (65), with resulting p-values reported. For boxplots shown in A.-F., H., J., and L., values between the 25^th^ (bottom) and 75^th^ (top) percentile are displayed, with the median reflected by the line within; whiskers reach the minimum and maximum values.

To understand the metabolic consequences of *Pptc7* loss in adult animals, as well as the potential origins of their lower body mass, we performed indirect calorimetry using metabolic cages. As altered body mass can affect metabolic rates, we performed these studies on animals as early as 8 weeks post-*Pptc7* KO, which resulted in a cohort that had trending but non-significant decreases in body mass across sexes (Figures 1E, F). Interestingly, male–but not female–*Pptc7* KO animals had significantly decreased whole body oxygen consumption (i.e., VO_2_) across tested timepoints (Figures 1G, H, Supplemental Figures 1A, B). Consistently, male *Pptc7* KO animals had significantly lower averaged energy expenditure per hour (Figures 1I, J), resulting in lower cumulative energy expenditure over a two-day period (Figure 1K). Female *Pptc7* KO mice did not show significant differences in energy expenditure across tested timepoints (Supplemental Figures 1C, D), nor did either sex display alterations in locomotor activity (Supplemental Figures 1E-F). Interestingly, only female *Pptc7* KO animals showed significant or trending decreases in respiratory exchange ratio (Supplemental Figure 1G-H), further indicating sex-specific metabolic alterations upon loss of PPTC7 in mice.

As energy expenditure and body mass are influenced by food consumption, we analyzed energy intake (i.e., food mass consumed x caloric content) across animal cohorts. Though neither sex of *Pptc7* KO animals displayed significant differences in averaged energy intake per hour (Figure 1L, Supplemental Figure 1I), an analysis of cumulative energy intake revealed that both sexes of *Pptc7* KO animals consumed less food than their littermate controls over time (Figure 1M, Supplemental Figure 1J). Consistently, both sexes of *Pptc7* KO animals had a lower cumulative energy balance than their floxed littermates (Figure 1N, Supplemental Figure 1K), likely contributing to their decreased body mass.

The trending lower weight of *Pptc7* KO animals in our cohorts may influence organismal metabolism, as smaller animals typically require less energy intake and respire less than larger animals (*21*). To distinguish contributions of body weight versus *Pptc7* KO-specific effects to the phenotypes observed in these experiments, we performed ANCOVA to statistically evaluate difference across genotypes (e.g., floxed and *Pptc7* KO animals) for mass-dependent parameters such as oxygen consumption rates, energy expenditure, and energy intake. If decreases in these phenotypes stem solely from mass, datapoints should lie on the same regression line, irrespective of genotype. However, if data segregate into lines that are shifted from one another, this suggests genotype-dependent contributions to phenotypes (i.e., group effects). ANCOVA analysis revealed that, in male *Pptc7* KO mice, VO_2_ and energy expenditure displayed a significant group effect, with a trending but non-significant effect from mass (Figures 1O, 1P). On the contrary, food intake showed a significant mass effect independent of group effect (Figure 1Q). These analyses suggest that the diminished food intake in male *Pptc7* KO animals derives largely from their smaller mass, whereas the decreases in oxygen consumption and energy expenditure result from genotype-specific effects due to loss of *Pptc7*. Like male mice, female animals showed a significant mass-dependent decrease in cumulative energy intake, but neither oxygen consumption rates nor energy expenditure reached significance for group-dependent (e.g., *Pptc7* KO-driven) effects (Supplemental Figures 1L-N). Collectively, these data demonstrate that both female and male *Pptc7* KO animals have diminished body weight and lean mass over time, but only males manifest significant and consistent decreases in respiratory capacity and energy expenditure due to loss of *Pptc7*.

### PPTC7 KO animals have altered circulating metabolites suggesting systemic mitochondrial dysfunction

Loss of *Pptc7* disrupts various metabolic pathways in cultured cells (*7*, *19*, *22*), perinatal mice (*6*), and human patients (*19*), but whether similar dysfunction manifests in adult animals has not been tested. Given the alterations in energy expenditure in *Pptc7* KO male mice at 20 weeks post-tamoxifen treatment, we analyzed circulating metabolites in animals at this timepoint. We collected plasma and detected 167 unique polar metabolites via LC-MS, forty of which were significantly altered in *Pptc7* KO animals relative to floxed controls (Figure 2A).

**Figure 2:**
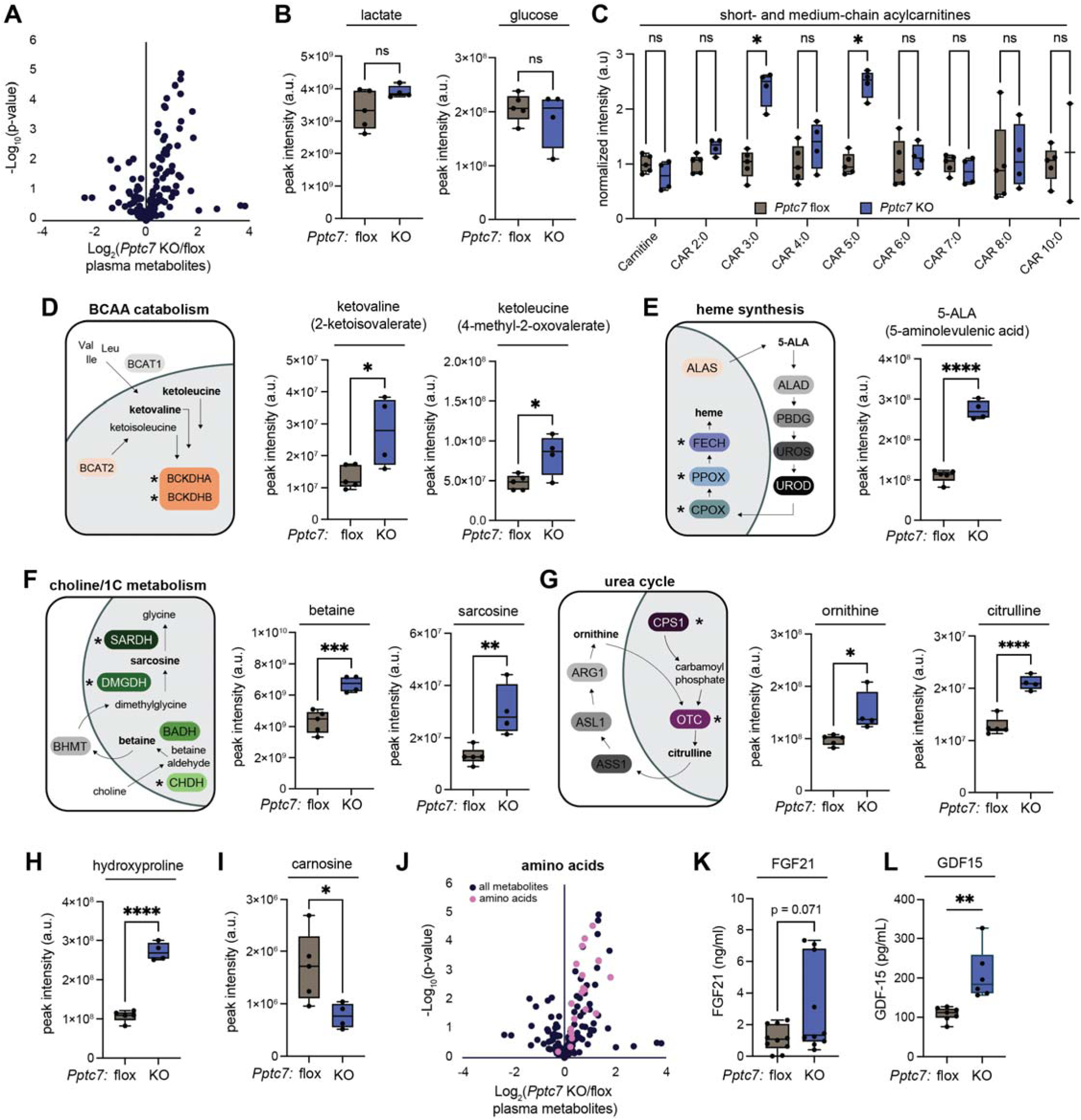
PPTC7 knockout animals have altered circulating metabolites suggesting systemic mitochondrial dysfunction. **A**. Volcano plot displaying 167 polar metabolites detected in plasma isolated from *Pptc7* flox (n=5) or *Pptc7* KO (n=4) male animals aged 20 weeks post-tamoxifen treatment. Statistical analysis performed using an unpaired Student’s t-test. **B**. Lactate and glucose levels as quantified by integrated peak area from LC-MS in plasma from control and KO animals. n.s. = not significant, Welch’s t-test. **C**. Short- and medium-chain acylcarnitines found in circulating plasma of *Pptc7* flox and KO animals. * = p<0.05, Lognormal t test via the Holm-Šídák method. **D**.-**G**. Metabolite pathways with significant changes in *Pptc7* KO animals. For each diagram, proteins affected in liver are shown in color and with an asterisk to indicate their downregulation; primary data associated with each protein annotated can be found in Supplemental Figure 2. D. Branched chain amino acid (BCAA) catabolites as quantified by integrated peak area from LC-MS in plasma from control and KO animals. * = p<0.05, Welch’s t-test. E. Heme metabolites as quantified by integrated peak area from LC-MS in plasma from control and KO animals. **** = p<0.0001, Welch’s t-test. F. Choline/one carbon (1C) metabolites as quantified by integrated peak area from LC-MS in plasma from control and KO animals. ** = p<0.01, *** = p<0.001, Welch’s t-test. G. Urea cycle metabolites as quantified by integrated peak area from LC-MS in plasma from control and KO animals. * = p<0.05, **** = p<0.0001, Welch’s t-test. **H**., **I**. Hydroxyproline (H.), carnosine (I.) as quantified by integrated peak area from LC-MS in plasma from control and KO animals. * = p<0.05, **** = p<0.0001, Welch’s t-test. **J**. Volcano plot highlighting amino acids quantified in *Pptc7* floxed versus KO animals. Statistical analysis performed using an unpaired Student’s t-test. **K**., **L**. FGF21 (K.) and GDF15 (L.) levels quantified from circulating serum in overnight fasted male mice aged 20 weeks post-tamoxifen treatment. ** = p<0.01, Welch’s t-test. For all boxplots shown, values between the 25^th^ (bottom) and 75^th^ (top) percentile are displayed, with the median reflected by the line within; whiskers reach the minimum and maximum values.

We first compared metabolite profiles from adult *Pptc7* KO mice to our previous datasets collected in perinatal *Pptc7* KO animals (*6*). Perinatal *Pptc7* KO animals presented with hypoketotic hypoglycemia, elevated circulating lactate, and increased tissue-resident acylcarnitines (*6*), indicative of defects in fatty acid oxidation (FAO) (*23*). Interestingly, adult *Pptc7* KO animals failed to display similar metabolic dysfunction, with no significant differences seen in circulating lactate or glucose (Figure 2B), nor in most quantified short- and medium-chain acylcarnitines (Figure 2C). Two notable exceptions were C3:0 carnitine (i.e., propionoylcarnitine) and C5:0 carnitine (i.e, valerylcarnitine), which each showed significant elevation in adult *Pptc7* KO animals (Figure 2C). These acylcarnitine species derive from the breakdown of odd-chain fatty acids or via the catabolism of the branched chain amino acids (BCAAs). Given the lack of elevation in C7:0 acylcarnitine (which would be expected if a defect in odd chain FAO were present), our data suggest that the elevated C3:0 and C5:0 acylcarnitines likely arise due to compromised BCAA catabolism. Consistently, α-ketoisovalerate (i.e., ketovaline) and 4-methyl-2-oxovalerate (i.e., ketoleucine), were significantly elevated in *Pptc7* KO plasma (Figure 2D), suggesting loss of PPTC7 promotes defects at multiple points within the BCAA catabolic pathway. Notably, significant elevation of propionoylcarnitine and ketoleucine were also found in plasma from human patients harboring mutations in *PPTC7* (*19*), as well as in liver and heart tissue in the perinatal *Pptc7* KO animals (*6*). Collectively, these data suggest that PPTC7 expression enables FAO in select developmental contexts, but loss of *Pptc7* compromises BCAA catabolism across multiple mouse models as well as in human patients.

Examination of other changing polar metabolites further suggested multi-systemic metabolic dysfunction due to loss of PPTC7. Amongst the most significantly altered metabolites in *Pptc7* knockout animals was aminolevulinic acid, or ALA, the first precursor of heme biosynthesis (Figure 2E). Abnormal circulating ALA is associated with porphyria, which originates in the bone marrow (as erythropoetic porphyria) or the liver (as hepatic porphyria) (*24*). *Pptc7* KO mice display elevated levels of other metabolites that may stem from hepatic dysfunction, including betaine and sarcosine (Figure 2F), intermediates in choline metabolism, and citrulline and ornithine (Figure 2G), two metabolites within the urea cycle. We previously performed proteomics on liver tissue from inducible *Pptc7* KO animals two weeks after tamoxifen-induced knockout (*7*), which revealed significant downregulation of many mitochondrial proteins, including components of the BCAA dehydrogenase complex (i.e., BCKDHA, BCKDHB, Supplemental Figure 2A), sarcosine dehydrogenase (SARDH, Supplemental Figure 2B), urea cycle components (e.g., ornithine carbamoyltransferase (OTC) and carbamoyl phosphate synthase (CPS1), Supplemental Figure 2C) and later enzymes within the heme biosynthetic pathway (e.g., Protoporphyrinogen oxidase (PPOX) and ferrochelatase (FECH), Supplemental Figure 2D). These data suggest that altered circulating metabolites found in UBC-Cre *Pptc7* knockout animals likely accumulate due to decreased expression of catabolic enzymes within hepatic mitochondria.

Other circulating catabolites found in adult inducible *Pptc7* KO mice suggested dysfunction in connective tissue and skeletal muscle. The most significantly elevated circulating metabolite identified in our analysis was hydroxyproline (Figure 2H)–a modified amino acid derivative enriched in collagen that is released into circulation upon its degradation. Notably, circulating carnosine was significantly decreased in *Pptc7* KO animals (Figure 2I); previous studies have associated lower bloodborne carnosine with muscle wasting and cachexia in cancer patients (*25*). Furthermore, grouping classes of metabolites revealed a stark trend for circulating amino acids, which were almost all elevated in the plasma of *Pptc7* KO mice relative to their control littermates (Figure 2J). Given the importance of skeletal muscle as a depot for systematic amino acid availability, coupled with our data suggesting systemic mitochondrial dysfunction, we considered that *Pptc7* KO animals may also show elevated levels of FGF-21 and GDF-15, two circulating biomarkers previously associated with mitochondrial myopathies (*26*). We collected serum from male floxed or *Pptc7* knockout animals 20-weeks post-tamoxifen that were fasted overnight and found trending or significant elevation of circulating FGF-21 and GDF-15 (Figures 2K, L). Given the elevation of metabolites and hormones previously associated with skeletal muscle dysfunction, coupled with our data that *Pptc7* KO animals manifest lower lean mass and body weight, we explored the possibility that these animals manifest skeletal muscle dysfunction.

### Loss of Pptc7 selectively impacts skeletal muscle mass

We first established that PPTC7 is expressed in skeletal muscle, with the GTEx database (*27*) demonstrating its relatively high gene expression amongst human tissues (Supplemental Figure 3A). We confirmed this at the protein level by western blotting PPTC7 across eight tissues from wild-type C57BL/6J mice. We found that, though expressed in all tested tissues, PPTC7 displayed higher protein levels in tissues of high energy demand such as the heart, skeletal muscle, kidney, and brain (Figures 3A, B). We validated tamoxifen-mediated excision in each of these tissues in *Pptc7* KO animals, noting robust KO in kidney and skeletal muscle, minor mosaicism in heart, and substantial mosaicism in the brain (Supplemental Figure 3B). Collectively, these data led us to hypothesize that the diminished lean mass and compromised oxygen consumption seen in *Pptc7* KO animals resulted from skeletal muscle dysfunction.

**Figure 3:**
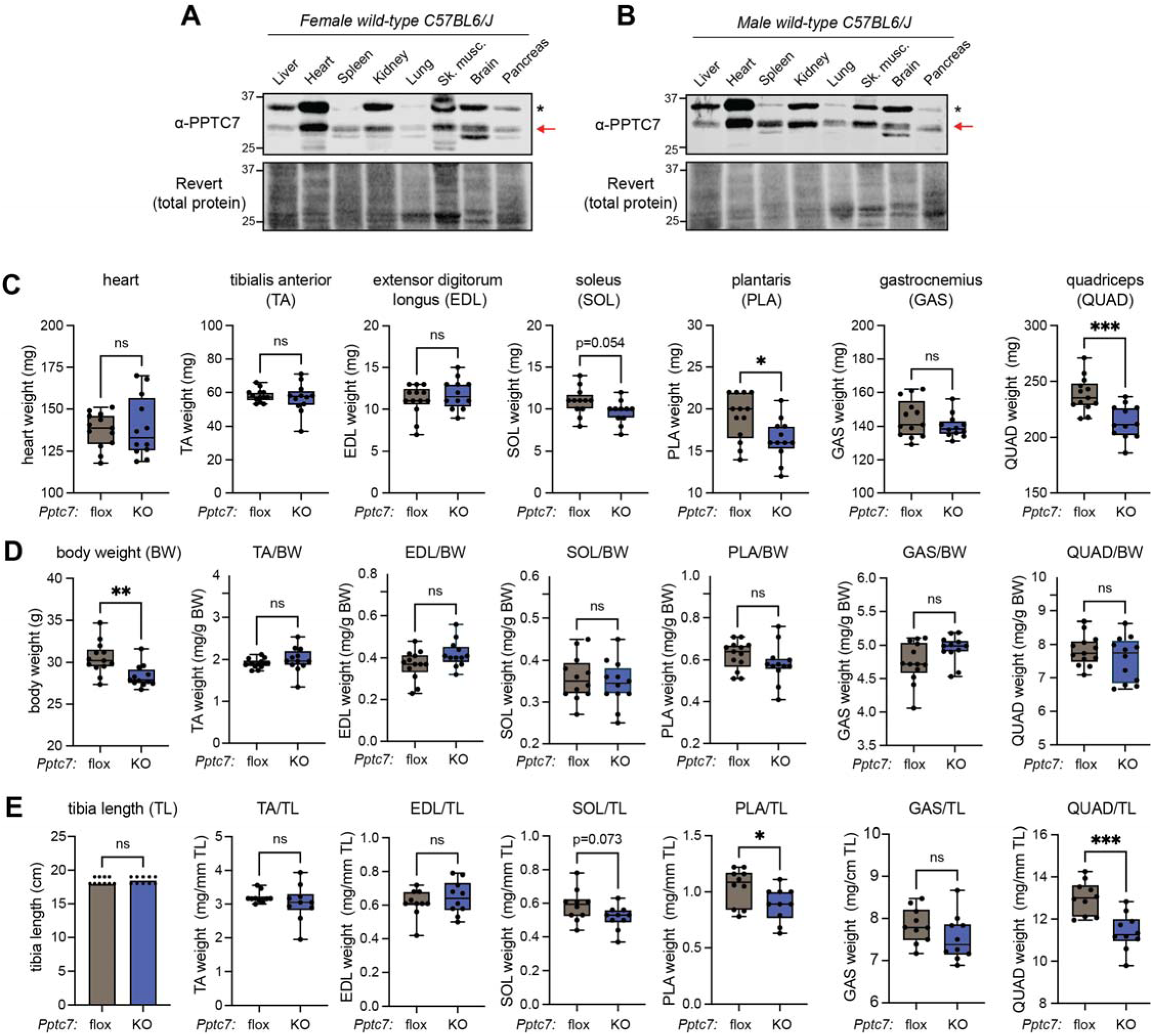
Loss of Pptc7 selectively diminishes skeletal muscle mass, contributing to decreased body weight. **A**., **B**. Western blot analysis of endogenous PPTC7 expression in a female (A.) or male (B.) wild-type C57BL6/J mouse. * = non-specific band, red arrow indicates PPTC7. Revert stain shown as a load control demonstrating total protein. **C**. Raw weights of various muscles isolated from *Pptc7* flox (n=13) or Pptc7 KO (n=12) male animals aged to 20 weeks post-tamoxifen treatment. n.s. = not significant, * = p<0.05, *** = p<0.001, Welch’s t-test. **D**. Body weight (left) and body weight (BW)-normalized muscle weights for *Pptc7* flox (n=13) or Pptc7 KO (n=12) male animals aged to 20 weeks post-tamoxifen treatment. n.s. = not significant, ** = p<0.01, Welch’s t-test. **E**. Tibia length (left) and tibia length (TL)-normalized muscle weights for *Pptc7* flox (n=13) or Pptc7 KO (n=12) male animals aged to 20 weeks post-tamoxifen treatment. n.s. = not significant, * = p<0.05, *** = p<0.001, Welch’s t-test. For all boxplots shown, values between the 25^th^ (bottom) and 75^th^ (top) percentile are displayed, with the median reflected by the line within; whiskers reach the minimum and maximum values.

We aged control and *Pptc7* KO animals of both sexes to 20 weeks post-tamoxifen treatment and measured the weights of six skeletal muscles of the hindlimb. We selected four extensor muscles that weight bear during locomotion (the gastrocnemius (GAS), plantaris (PLA), soleus (SOL), and quadriceps (QUAD)) and two extensor muscles that do not weight bear during locomotion (the tibialis anterior (TA) and extensor digitorum longus (EDL)). In addition to these functional differences, these muscles provide a distribution of slow oxidative (i.e., type I), fast oxidative (i.e., type IIA) and fast glycolytic (i.e., types IIB and IIX) fiber types (Figure 3C). Despite presenting with decreased lean mass and body weight, we found no significant differences in the masses of tested hindlimb muscles from female *Pptc7* knockout mice (Supplemental Figure 3C). However, muscles from male *Pptc7* knockout mice showed lower mass, as the PLA and QUAD muscles weighed significantly less than matched muscles from *Pptc7* floxed animals (Figure 3C), and the SOL trended as smaller (p=0.054, Figure 3C). Interestingly, this decrease in mass was not evident across muscles, as the TA, EDL, and GAS displayed non-significant trends in size (Figure 3C). Furthermore, the heart, another striated muscle type, did not differ in mass between *Pptc7* KO and control animals (Figure 3C). These data suggest that chronic (i.e., 20-week), tamoxifen-induced loss of *Pptc7* causes reduced mass of select muscles in a sex-specific manner.

We considered that the observed decreases in skeletal muscle weight may stem from altered body composition or decreases in the skeletal size of animals. We sought to distinguish these possibilities by normalizing muscle masses to body weight (as a proxy for body composition) or tibia length (as a proxy for skeletal size). Like our initial cohort, male *Pptc7* knockout animals weighed significantly less than control floxed littermates (Figure 3D), yet showed comparable tibia length (Figure 3E), suggesting the decreased lean mass of male *Pptc7* KO animals drives their lower body weights independent of changes in linear size. Consistently, no differences were observed between *Pptc7* knockout and control muscles when normalized to body weight (Figure 3D), whereas normalization of *Pptc7* KO muscle weights to tibia length did not alter the trends seen across raw muscle weights (Figure 3E). These data suggest that the loss of lean mass at least partially stems from lowered skeletal muscle mass, and that this loss in muscle mass is proportional to the body weight rather than body length of male *Pptc7* knockout animals.

### Pptc7 KO skeletal muscles show selective decreases in fiber size and variable patterns of fiber type switching

To test the effects of *Pptc7* KO at the fiber level, we cryopreserved the ankle plantarflexor (GAS/PLA/SOL) muscle group and measured fiber area and fiber type distribution. We chose this muscle group as two of the muscles (SOL and PLA) showed differences in mass in male *Pptc7* knockout mice, and because these muscles include all four murine fiber-types, allowing us to test whether effects were fiber-type specific. As muscles can also undergo fiber-type switching, these experiments also allowed us to quantify the fiber-type distribution across genotypes. We isolated and sectioned muscle bundles from control and *Pptc7* KO mice and used a cocktail of myosin heavy chain isoform-specific antibodies as well as a laminin antibody to visualize and quantify muscle fiber-types using fluorescence microscopy. Representative images of control and *Pptc7* KO animals are shown for the soleus (SOL, primarily type I and type IIA fibers, Figure 4A), the plantaris (PLA, primarily types IIA, IIB, IIX fibers, Figure 4C), the red gastrocnemius (R-GAS, all fiber types present, Figure 4E), and the white gastrocnemius (W-GAS, primarily type IIB fibers, Figure 4G).

**Figure 4:**
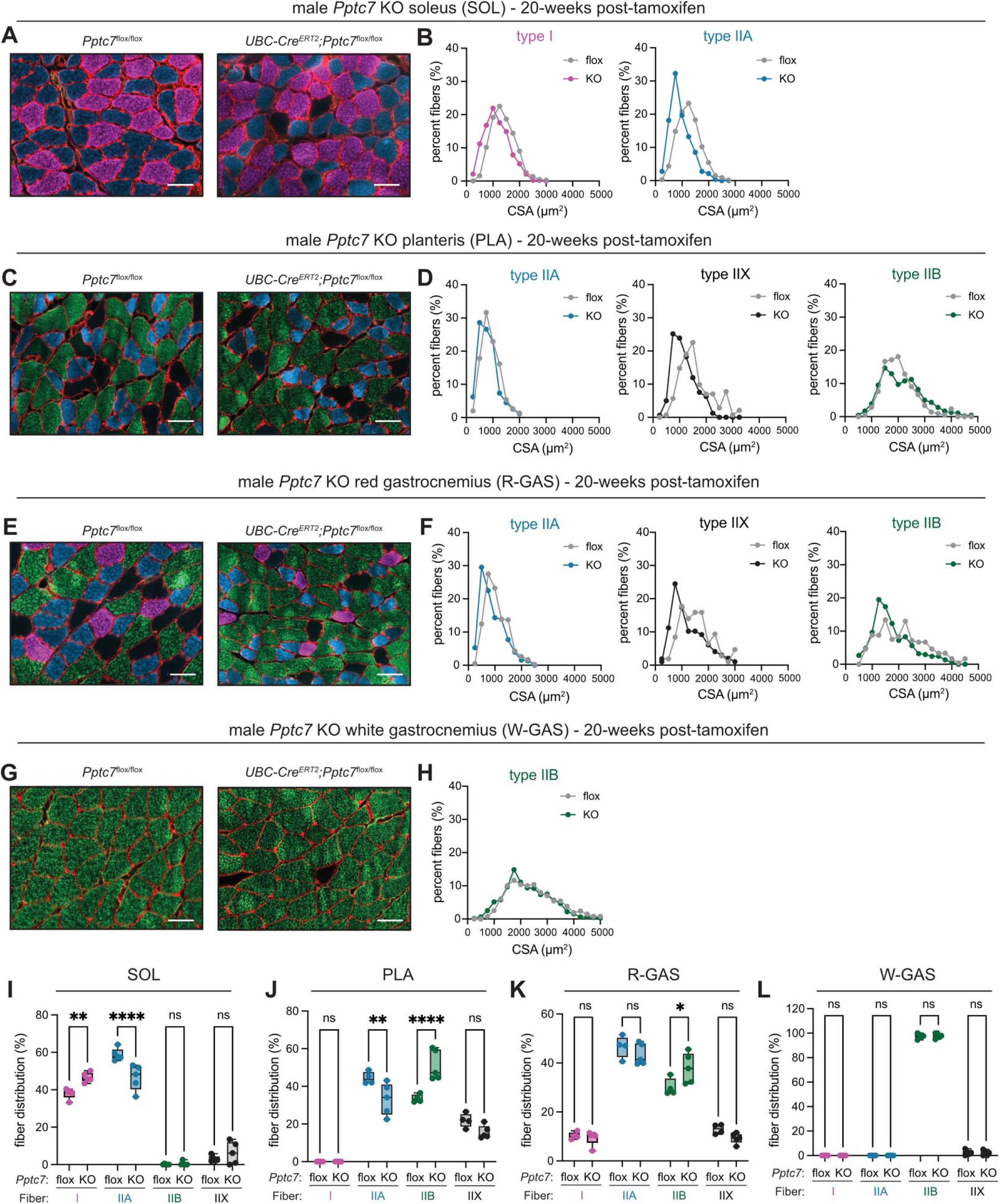
Pptc7 KO skeletal muscles show selective decreases in fiber size and variable patterns of fiber type switching. **A**. Representative images of soleus (SOL) from *Pptc7* flox (left) and *Pptc7* KO (right) animals. Scale bar = 50 μm. **B**. Histogram distribution of fibers quantified from *Pptc7* flox and *Pptc7* KO male mice aged to 20 weeks post-tamoxifen treatment. **C**. Representative images of plantaris (PLA) from *Pptc7* flox (left) and *Pptc7* KO (right) animals. Scale bar = 50 μm. **D**. Histogram distribution of fibers quantified from *Pptc7* flox and *Pptc7* KO male mice aged to 20 weeks post-tamoxifen treatment. **E**. Representative images of red gastrocnemius (R-GAS) from *Pptc7* flox (left) and *Pptc7* KO (right) animals. Scale bar = 50 μm. **F**. Histogram distribution of fibers quantified from *Pptc7* flox and *Pptc7* KO male mice aged to 20 weeks post-tamoxifen treatment. **G**. Representative images of white gastrocnemius (W-GAS) from *Pptc7* flox (left) and *Pptc7* KO (right) animals. Scale bar = 50 μm. **H**. Histogram distribution of fibers quantified from *Pptc7* flox and *Pptc7* KO male mice aged to 20 weeks post-tamoxifen treatment. **I**.-**L**. Fiber type distribution of type I (pink), type IIA (blue), type IIB (green), and type IIX (black) in SOL (I.), PLA (J.), R-GAS (K.) and W-GAS (L.). n.s. = not significant, * = p<0.05, ** = p<0.01, *** = p<0.001, ****=p<0.0001, Ordinary Two-way ANOVA. For all boxplots shown, values between the 25^th^ (bottom) and 75^th^ (top) percentile are displayed, with the median reflected by the line within; whiskers reach the minimum and maximum values.

We first analyzed fiber CSA, hypothesizing that the SOL and PLA muscles in *Pptc7* KO male mice would have smaller fibers, consistent with their relatively lower mass. Indeed, the SOL from *Pptc7* KO animals displayed a left-shifted distribution of type I and type IIA fiber size (Figure 4B), with both fiber types significantly smaller in *Pptc7* knockout SOL relative to floxed controls (Supplemental Figure 4A). Interestingly, different trends were seen for the PLA, as type IIA and type IIX fibers displayed left-shifted fiber size distribution and significantly decreased CSA (Figure 4D, Supplemental Figure 4B), whereas type IIB fibers showed no difference (Figure 4D, Supplemental Figure 4B). Interestingly, all type II fiber types in R-GAS displayed a left-shifted distribution in *Pptc7* KO animals, but type I fiber CSA was not significantly altered (Figures 4F, Supplemental Figures 4C, D). Similarly, W-GAS had a slight but significant shift in type IIB fiber CSA in *Pptc7* KO animals relative to control littermates. (Figures 4G, 4H, Supplemental Figure 4E).

We also sought to determine whether fiber-type switching occurred in *Pptc7* KO animals, and quantified the relative distribution of type I, type IIA, type IIB, and type IIX fiber-type frequencies across each muscle in control and *Pptc7* KO animals (Figure 4I-L). These data revealed that fiber type switching was evident in most muscles examined, but was neither uniform in its directionality (e.g., oxidative to glycolytic) nor in the specific fiber types affected. The SOL muscle from *Pptc7* KO animals displayed significantly fewer type IIA fibers but significantly more type I fibers than their floxed littermates, undergoing an oxidative shift (Figure 4I). On the contrary, the PLA becomes more glycolytic with a shift from type IIA fibers to type IIB fibers (Figure 4J). The R-GAS of *Pptc7* KO animals displayed a significant increase in the percentage of type IIB fibers, but only trended towards lower type IIA and IIX fibers (Figure 4K). Finally, there was no evidence of a change in fiber-type distribution in the W-GAS of *Pptc7* KO mice (Figure 4L). These data suggest that sustained loss of *Pptc7* induces variable decreases in size and fiber distribution across hindlimb skeletal muscles.

### BNIP3 expression is increased across Pptc7 KO skeletal muscles

The variability in responses to *Pptc7* KO across skeletal muscles led us to explore potential mechanisms giving rise to such diverse phenotypes. As PPTC7 has recently been established to diminish receptor-mediated mitophagy by limiting the expression of BNIP3 and NIX (*7–10*), we considered that dysregulation or one or both of these receptors may underlie the dysfunction seen in *Pptc7* KO skeletal muscle. Though each of these mitophagy receptors have been linked to skeletal muscle biology (*28–32*), GTEx data (*27*) demonstrate that BNIP3 has higher expression than NIX in human skeletal muscle (Supplemental Figures 5A, B), which we confirmed via western blotting across tissues in wild-type, C57BL6/J mice (Figures 5A, B, Supplemental Figures 5C, D). These data prompted us to focus on BNIP3, hypothesizing that its differential upregulation may drive the variable dysfunction seen across hindlimb skeletal muscles in *Pptc7* KO animals.

**Figure 5:**
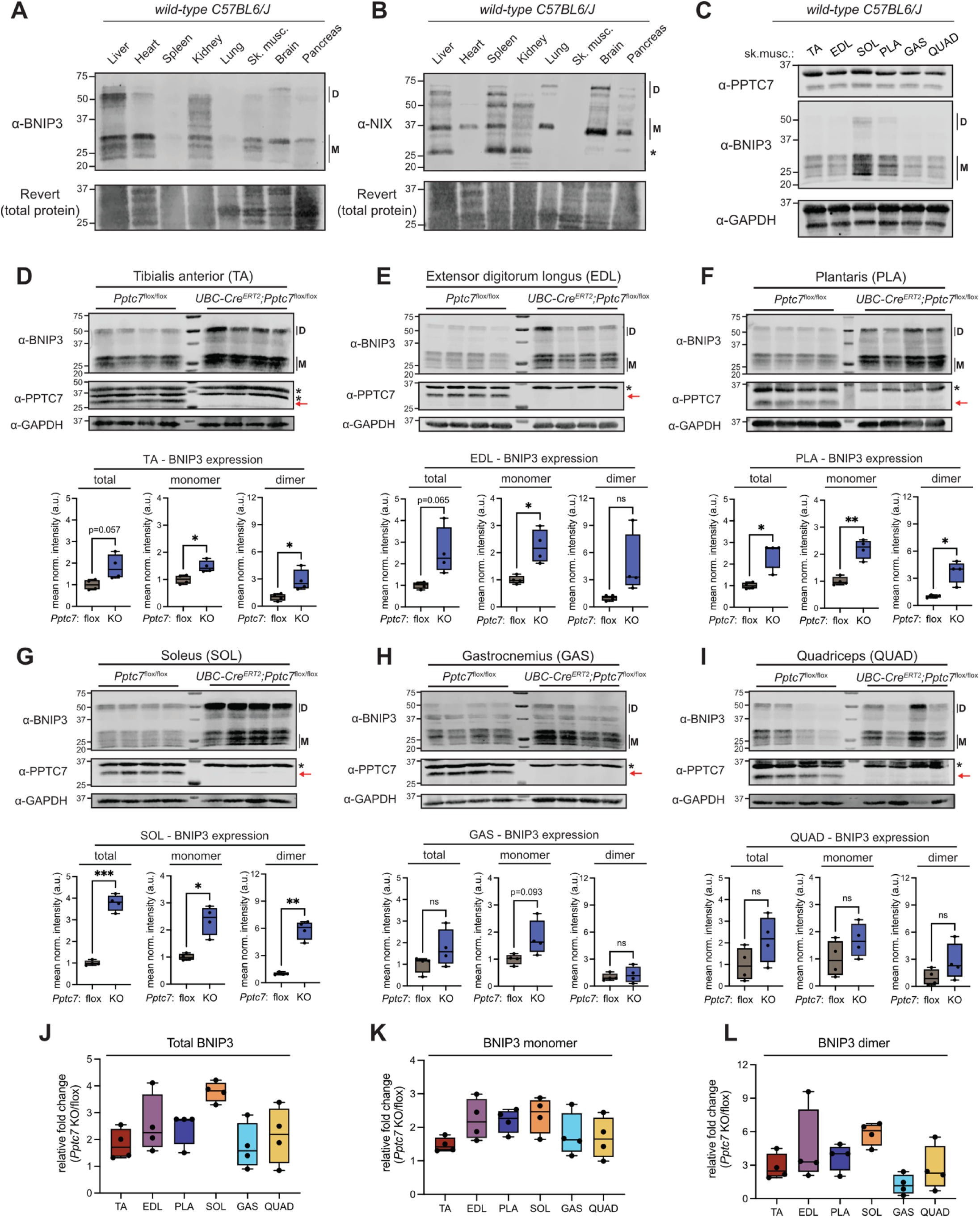
BNIP3 expression is increased across Pptc7 KO skeletal muscles. **A., B**. BNIP3 (A.) and NIX (B.) total protein levels across representative tissues from a wild-type C57BL6/J mouse as assayed by western blotting. D = dimeric form, M = monomeric form. * = non-specific band. Revert total protein stain shown as loading control. **C**. PPTC7 (top) and BNIP3 (middle) total protein levels across six different skeletal muscles from a wild-type C57BL6/J mouse as assayed by western blotting. GAPDH blot shown as loading control. **D**.-**I**. Total BNIP3 (top) and PPTC7 (middle) levels as assayed by western blotting in TA (D.), EDL (E.), PLA (F.), SOL (G.), GAS (H.), and QUAD (I.) muscles from *Pptc7* flox (n=4) and *Pptc7* KO (n=4) male mice aged to 20 weeks post-tamoxifen. D = dimeric form, M = monomeric form. GAPDH blot shown as loading control. BNIP3 monomer and dimer populations were quantified and are shown as box plots below immunoblots. n.s. = not significant, * = p<0.05, ** = p<0.01, *** = p<0.001, ****=p<0.0001, Welch’s t test. **J**.-**L**. Relative quantification of ratio of BNIP3 levels in *Pptc7* KO/*Pptc7* flox animals for total BNIP3 (J.), BNIP3 monomer (K.) and BNIP3 dimer (L.) across six different skeletal muscles. For all boxplots shown, values between the 25^th^ (bottom) and 75^th^ (top) percentile are displayed, with the median reflected by the line within; whiskers reach the minimum and maximum values.

We first considered that the six tested hindlimb skeletal muscles may express different levels of BNIP3 or PPTC7 at steady state. We immunoblotted these targets in wild-type hindlimb muscle and found that while PPTC7 had uniform expression across tested muscles, BNIP3 levels differed across muscle types (Figure 5C), with highest levels in the SOL. Though interesting, no obvious trend in basal BNIP3 protein levels emerged to explain the differential mass seen in *Pptc7* KO skeletal muscles. This led us to hypothesize that BNIP3 may be upregulated to different extents upon loss of *Pptc7* across hindlimb muscles. We generated lysates from control or *Pptc7* KO animals from the six selected hindlimb muscles and performed quantitative western blotting for BNIP3 (Figures 5D-I). Notably, our gel resolving conditions allowed the resolution of monomeric and dimeric BNIP3 species, and, as dimers forms are thought to be the more active forms of these receptors (*33*, *34*), we quantified the upregulation of each isoform as well as total BNIP3, which revealed unanticipated trends. First, the TA and EDL displayed significant upregulation of at least one BNIP3 species in *Pptc7* KO animals (Figures 5D, E) despite neither of these muscles displaying altered mass. Second, the PLA and SOL muscles displayed significant upregulation of all quantified BNIP3 species in *Pptc7* KO animals (Figures 5F, G), despite presenting with variable fiber type switching, as *Pptc7* KO PLA showed a glycolytic shift (type IIA to type IIB fibers) whereas *Pptc7* KO SOL displayed an oxidative shift (type IIA to type I fibers). Finally, the GAS and QUAD muscles, which displayed significant signs of dysfunction by at least one metric (decreased CSA or mass, respectively), showed insignificant and variable trends in BNIP3 protein levels between control and *Pptc7* KO animals (Figure 5H, I). Comparison between the fold changes of total, monomeric, and dimeric forms of BNIP3 demonstrated that the SOL had the highest upregulation of BNIP3 across all *Pptc7* KO muscle types (Figure 5J), largely driven by increased levels of the dimer species (Figure 5L), as monomeric species of BNIP3 are similar across muscle types (Figure 5K). Importantly, this variability in BNIP3 upregulation does not seem to be caused by incomplete perturbation of *Pptc7*, as immunoblots revealed its robust loss at the protein level across all six tested muscle types (Figures 5D-I). Instead, these data suggest that genetic perturbation of *Pptc7* induces surprisingly variable patterns of BNIP3 upregulation across skeletal muscle types.

### Pptc7 KO muscles have relatively normal mitochondrial form and function but show variable losses in mitochondrial content

The upregulation of BNIP3 across *Pptc7* KO muscles did not fully explain the differential dysfunction across skeletal muscles and led us to explore mitochondrial form and function across these tissues. We first examined the SOL and EDL muscles, as each muscle displayed significant upregulation of at least one form of BNIP3 (Figures 5E, F) but differential susceptibility to diminished mass in *Pptc7* KO animals (Figure 3C). Furthermore, the SOL is a mitochondria-rich oxidative muscle while the EDL is glycolytic, leading us to hypothesize that the SOL muscle would manifest greater levels of mitochondrial dysfunction than the EDL in *Pptc7* KO animals.

To examine mitochondrial ultrastructure, we performed transmission electron microscopy on muscles isolated from control and *Pptc7* KO mice aged to 20-weeks post-tamoxifen treatment. We noted at least two different morphologies of mitochondria in the SOL muscles of control and *Pptc7* KO mice. The first, which appeared to be interfibrillar mitochondria (IFM), were smaller and more sparsely populated across the muscle (Figures 6A, B, left panels), which may reflect mitochondria found in type IIA skeletal muscle fibers. The second mitochondrial population found in the SOL muscles of both genotypes were larger, more cristae-dense, and found in the interfibrillar region (Figures 6A, B, middle and right panels). Though we noted some differences in density and intensity of the counterstaining across genotypes, the ultrastructure of mitochondria was relatively normal in floxed and *Pptc7* KO animals. However, one striking phenotype in the *Pptc7* KO SOL muscle was an increased frequency of low-density regions that contained whorls of membrane or cellular structures and likely represent autophagosomes (Figure 6B, lower panels). Examination of the EDL revealed similar trends, with similar mitochondrial ultrastructure across control and *Pptc7* KO tissues but with a notable increase in autophagosome-like structures in *Pptc7* KO muscle (Figures 6C, D). Quantification revealed a significant increase in autophagosomes in both *Pptc7* KO muscle types (Figures 6E, F), though the increase is larger in the SOL than the EDL. Collectively, these data suggest that the SOL and EDL undergo elevated autophagy, and, in light of their upregulation of BNIP3, this likely reflects increased mitophagy.

**Figure 6:**
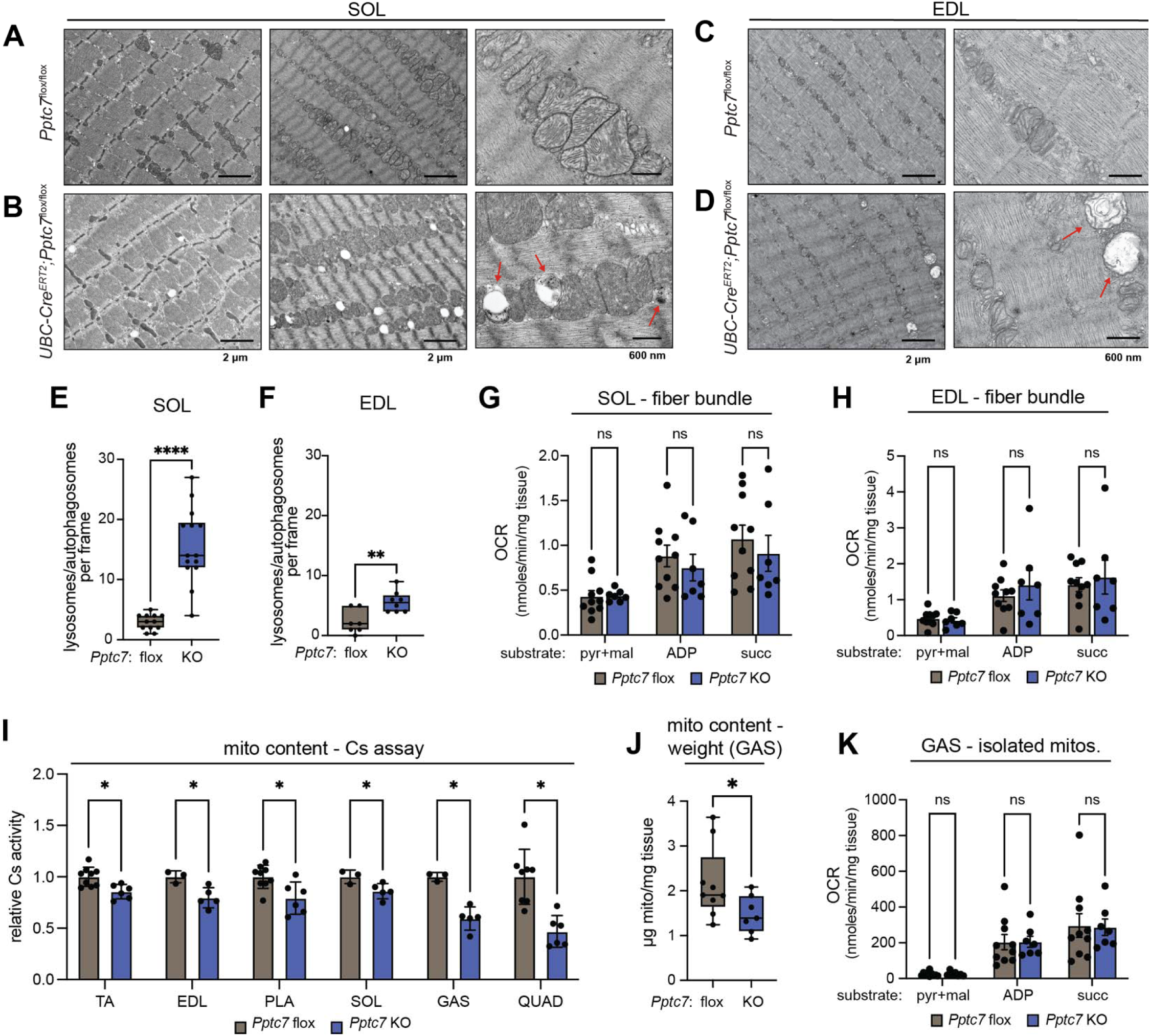
Pptc7 knockout muscles have relatively normal mitochondrial form and function but show variable losses in mitochondrial content.. **A.-D**. Transmission electron micrographs (TEM) for control (top) or experimental (bottom) SOL (A., B.) and EDL (C.,D.) muscle tissue. Scale bars are indicated below images for associated column. Red arrows indicate autophagosome-like structures. **E**., **F**. Quantification of autophagosomes from TEM images for SOL (E.) and EDL (F.). ** = p<0.01, ****=p<0.0001, Welch’s t test. **G**., **H**. Oxygen consumption rates (OCR) in permeabilized muscle fibers from the SOL (G.) or EDL (H.) muscles. Data analyzed with multiple unpaired t tests using the Holm-Šídák method for multiple comparisons. n.s. = not significant. **I**. Citrate synthase assays from muscle lysates of corresponding tissues. Data analyzed with multiple unpaired t tests using the Holm-Šídák method for multiple comparisons. * = p<0.05. **J**. Isolated mitochondrial content normalized to to milligrams of wet weight tissue. * = p<0.05, ****=p<0.0001, Welch’s t test. **K**. Oxygen consumption rates (OCR) of isolated mitochondria normalized for total mitochondrial content. Data analyzed with multiple unpaired t tests using the Holm-Šídák method for multiple comparisons. n.s. = not significant. For all boxplots shown, values between the 25^th^ (bottom) and 75^th^ (top) percentile are displayed, with the median reflected by the line within; whiskers reach the minimum and maximum values

To assess mitochondrial function, we performed Oroboros-based respirometry on permeabilized muscle fibers from control and *Pptc7* KO male mice. We hypothesized that *Pptc7* KO muscles would have diminished mitochondrial respiration due to decreased mitochondrial content caused by elevated BNIP3-mediated mitophagy. However, we found no significant differences in respiratory capacity between the SOL and EDL muscles from control and *Pptc7* KO animals (Figures 6G, H). We then sought to quantify mitochondrial content by performing citrate synthase (Cs) activity assays from lysates derived from the TA, EDL, PLA, SOL, GAS, and QUAD muscles from male *Pptc7* KO animals and their floxed littermates. Though all *Pptc7* KO muscles harbored significant decreases in Cs activity relative to control muscles, these experiments again revealed high variability across muscles (Figure 6I). The SOL and TA muscles displayed mild though significant decreases in content, each with a mean of ∼86% Cs activity relative to control muscles. The EDL and PLA showed slightly larger decreases in Cs activity, as *Pptc7* KO muscles had 79-80% signal relative to control muscles. These data may explain why we did not see robust oxygen consumption differences in *Pptc7* KO SOL or EDL muscle bundles, as the degree of diminished mitochondrial content in these muscles may be too small to be reproducibly detected through this method. The GAS and QUAD muscles showed more substantial decreases in Cs activity, with *Pptc7* KO GAS at 59% signal relative to control muscles, and *Pptc7* KO QUAD displaying an even larger decrease to 47%. These data suggest that though BNIP3 is upregulated across all *Pptc7* KO skeletal muscles, this manifests in variable degrees of mitochondrial loss, at least at the level of citrate synthase activity.

The larger decreases in Cs activity in the GAS and QUAD muscles presented an opportunity to examine whether the mitochondria in these muscles also retained functionality. If so, isolation of mitochondria from *Pptc7* KO GAS or QUAD muscles should reveal lower quantities of mitochondrial mass per wet weight tissue, which, when normalized for these content differences, should show equivalent respiratory rates to control tissues. Indeed, isolation of mitochondria from the GAS muscle revealed significantly lower yields from *Pptc7* KO tissue relative to those of floxed controls (Figure 6J), and, upon normalization, these mitochondria displayed no significant differences in respiratory capacity relative to control mitochondria (Figure 6K). Collectively, these data suggest that ablation of *Pptc7* in skeletal muscle decreases mitochondrial content, but organelles remain functional, at least in terms of respiratory capacity in a defined, in vitro system. However, the variability in decreased mitochondrial content across skeletal muscles likely contributes to the heterogeneous phenotypes seen across the six tested muscle types.

### Knockout of Bnip3 rescues decreased body weight, lean mass, and perinatal lethality in Pptc7 knockout animals

The upregulation of BNIP3 across skeletal muscles and our data suggesting that mitochondria are largely functional in *Pptc7* KO skeletal muscles led us to hypothesize that excessive BNIP3-mediated mitophagy contributes to the pathologies seen in *Pptc7* KO animals. We generated two independent *Pptc7*/*Bnip3* double knockout animal models to test this. First, we made inducible *Pptc7*/*Bnip3* double knockout mice (herein referred to as inducible DKO mice) by crossing the UBC-Cre-ER^T2^;*Pptc7*^flox/flox^ animals to a global, non-conditional *Bnip3* KO model (Figure 7A). All animals were bred to homozygosity for floxed *Pptc7* and *Bnip3* KO alleles, with experimental animals harboring the UBC-Cre transgene and control animals lacking this recombinase. We validated that, upon tamoxifen treatment, inducible DKO animals had ablated PPTC7 and BNIP3 protein expression via western blotting across seven tissues (Figure 7B). We then tested whether loss of *Bnip3* would suppress the age-dependent decreases in body weight and body mass seen in both sexes of the inducible *Pptc7* KO animals. Following identical protocols, we tamoxifen treated DKO animals and aged them for 20-weeks post-tamoxifen treatment and analyzed trends in body weight and body composition using EchoMRI. Comparing these DKO animals to our initial cohorts, we found that both female and male DKO mice had increased body weight and significantly increased lean mass relative to inducible *Pptc7* KO animals, with no significant trend in fat mass (Figures 7C, D). Collectively, these data suggest that *Pptc7* KO animals of both sexes have lower body weights and lean mass due to the dysregulation of BNIP3.

**Figure 7:**
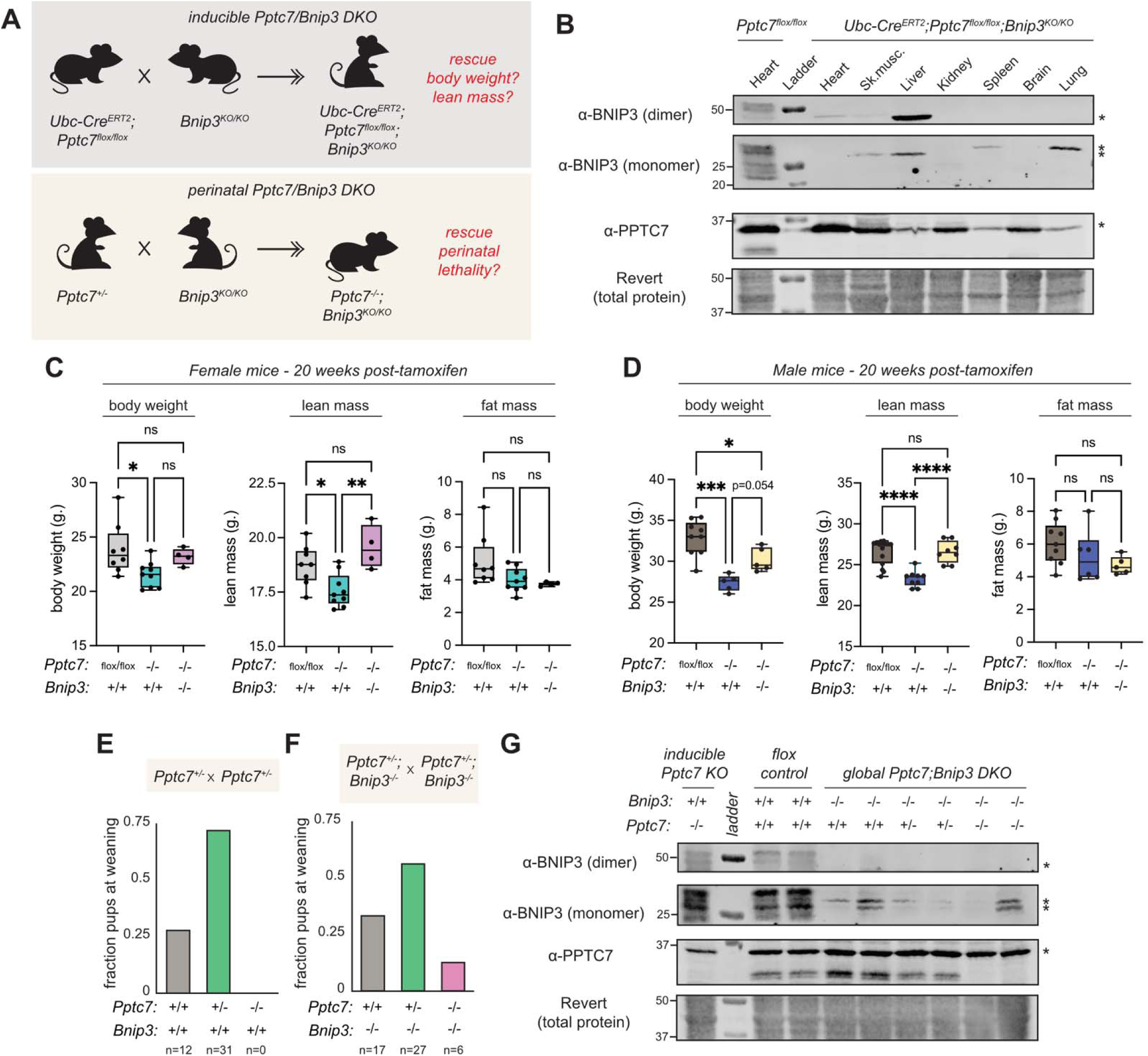
Knockout of Bnip3 rescues decreased body weight, lean mass, and perinatal lethality in Pptc7 knockout animals.. **A**. Graphic schematic of the generation of inducible *Pptc7*/*Bnip3* double knockout (DKO) mice (top) and global *Pptc7*/*Bnip3* DKO mice (bottom). **B**. Western blot of tissues from inducible *Pptc7*/*Bnip3* double mice. One Pptc7flox/flox heart tissue was used as a positive control left of the ladder. Revert total protein stain shown as a load control. * = non-specific band. **C**., **D**. Total body weight (left), lean mass (center) and fat mass (right) for control (*Pptc7* flox) and experimental (*Pptc7* KO and inducible *Pptc7*/*Bnip3* DKO) female (C.) and male (D.0 mice at 20 weeks post-tamoxifen treatment. n.s. = not significant, *=p<0.05, **=p<0.01, ***=p<0.001, ****=p<0.0001, Ordinary One-way ANOVA. E., F. Genotypes present at weaning (∼21 days of age) for crosses of *Pptc7*+/- animals (E.) and crosses of *Pptc7*+/-;*Bnip3*-/-DKO animals (F.). **G**. Western blot of liver tissue from global *Pptc7*/*Bnip3* double mice. UBC-Cre;Pptc7^flox/flox^ and Pptc7^flox/flox^ liver tissues were used as controls. Revert total protein stain shown as a load control.

Given the rescue in the inducible *Pptc7* KO model, we hypothesized that knockout of *Bnip3* may rescue other phenotypes, such as the perinatal lethality seen in the *Pptc7* global KO mice ((*6*), Figure 7E). We bred *Bnip3* global KO animals to our global, non-conditional *Pptc7* KO model, fixing the *Bnip3* KO alleles and interbreeding *Pptc7*+/- animals (Figure 7F). Though born at sub-Mendelian frequencies, 12.5% percent of animals across nine litters were null for *Pptc7* at weaning (Figure 7F), and full loss of PPTC7 and BNIP3 in global DKO animals was confirmed at the protein level via immunoblotting (Figure 7G). These data demonstrate that blunting BNIP3-mediated mitophagy can–but does not always–rescue perinatal lethality in the *Pptc7* KO background. The global *Pptc7*/*Bnip3* DKO animals that survived to weaning lived up to a year with no overt pathology, similar to global *Fbxl4* KO animals (*20*). Collectively, these rescue experiments suggest that excessive mitophagy at least partially drives perinatal lethality and decreased body weight in *Pptc7* KO models, underscoring the importance of PPTC7 maintenance of BNIP3-mediated mitophagy in vivo.

## Discussion

*Pptc7* is an essential gene for survival during the perinatal transition, as its germline knockout in mice causes fully penetrant death within a day of birth (*6*). Such a stark phenotype suggested that PPTC7 may promote mitochondrial metabolism in contexts beyond the perinatal transition. Here, we find that loss of PPTC7 is compatible with viability during adulthood, yet is accompanied by organism-wide dysfunction, including diminished body weight, decreased lean mass, and sex-specific defects in respiratory capacity. One potential explanation for the discrepancy in phenotypes between the global and inducible *Pptc7* KO models could be that knockout of *Pptc7* disrupts distinct molecular pathways during the perinatal transition and adulthood. However, we previously performed proteomic analyses that indicated *Pptc7* knockout causes similar losses in mitochondrial protein content and elevated mitochondrial phosphorylation occupancies in liver tissue from both perinatal and adult animals (*7*). These data suggest that, despite inducing common molecular signatures, the loss of *Pptc7* is variably essential across developmental contexts. This developmentally selective essentiality is further supported by the perinatal lethal phenotype displayed by *Fbxl4* KO mice (*20*), which harbor a genetically distinct perturbation in the same pathway that suppresses BNIP3- and NIX-mediated mitophagy. These data, along with our data demonstrating that *Bnip3* genetic ablation can rescue *Pptc7* KO perinatal lethality, suggest that uncontrolled and excessive mitophagy is particularly detrimental in the hours after birth. Notably, other mouse models that disrupt tissue-wide autophagy (e.g., *Atg5* and *Atg7* KO) or its regulation (e.g., via Rag GTPase-dependent regulation of mTORC1) display similar manifestations of perinatal lethality (*35–37*). Intriguingly, the lifespan of these animal models can be extended through supplementation with exogenous nutrients, suggesting that an inability to fine-tune autophagic or mitophagic flux disrupts critical metabolic processes during this developmental transition. Further work will be needed to understand the extent to which BNIP3 and NIX-mediated mitophagy enable organismal metabolism in the perinatal period and beyond.

Despite the evidence that BNIP3- and NIX-mediated mitophagy is critical during the perinatal transition, much remains unknown about the consequences of their unchecked regulation in physiological contexts beyond this timeframe. Some clues came from the global *Fbxl4* KO model, as the handful of animals that survived the extrauterine transition were viable with minimal overt pathologies for at least 8-12 months of age (*20*). Interestingly, however, these surviving adult *Fbxl4* KO animals displayed decreased body mass relative to littermate controls (*20*), suggesting that dysregulated BNIP3- and NIX-mediated mitophagy may be sufficient to decrease body weight. Consistently, our data demonstrate that simultaneous genetic disruption of *Bnip3* in the inducible *Pptc7* KO model corrects decreases in body mass seen in *Pptc7* KO animals (*6*). Furthermore, evidence suggests that such body mass phenotypes may extend to humans carrying select *PPTC7* variants. We recently identified three patients with recessive, loss-of-function mutations in PPTC7, each of whom presented with profound hypotonia and low body weight (i.e., below 3^rd^ percentile) (*19*), suggesting skeletal muscle defects and decreased body mass are conserved responses to loss of PPTC7 function. Intriguingly, analysis of multiple genome-wide association studies (GWAS) via the Common Metabolism Diseases Knowledge Portal (CMDKP) revealed a “very strong” association (i.e., a HuGE score of 45) between *PPTC7* gene variants and body weight in humans (*38*, *39*). Collectively, these data suggest that proper regulation of BNIP3-mediated mitophagy plays a conserved role in maintaining body mass in mammals.

Interestingly, the decrease in body mass seen in inducible *Pptc7* KO animals resulted from diminished lean mass rather than changes in fat mass. Given that skeletal muscle makes up an estimated 30-40% of body mass in rodents (*40*) and PPTC7 is highly expressed in this tissue (*27*), we hypothesized that skeletal muscle dysfunction contributed to this differential in body mass. In support of this hypothesis, we found evidence for reductions in size in multiple muscles from the *Pptc7* KO hindlimb – either via lower muscle mass, lower fiber CSA, or both. Interestingly, however, not all muscles tested were equally affected. The ankle flexors (TA and EDL) had masses equivalent to littermates, while the ankle extensors (GAS, PLA and SOL) and the knee extensors (QUAD) had lower mass and/or fiber CSA which may arise from an interaction between *Pptc7* loss and muscle use. The ankle and knee extensors are weight-bearing during locomotion, while the ankle flexors are not, and their loading and metabolic needs are thought to be higher as a result. In models of hindlimb unloading, the ankle extensors decrease in size to a greater extent than the ankle flexors, an effect which is attributed to differences in use (*41–43*). However, mouse activity is unaffected by *Pptc7* loss (via beam breaks in metabolic cages), suggesting a downward pressure on muscle size distinct from unloading or disuse. Such pressures could dampen protein synthesis, increase protein degradation, or a combination of both. Though our data demonstrate that BNIP3 is elevated across *Pptc7* KO skeletal muscle, which likely elevates mitophagy and thus increases protein degradation, we cannot rule out contributions of altered protein synthesis to the phenotypes described, particularly as mice were still growing during the post-tamoxifen period.

Differences in fiber type composition could also contribute to the differential response of the hindlimb muscles to *Pptc7* KO. The SOL is the slowest muscle and the only muscle examined with more than a few percent of oxidative type I fibers. The QUAD, GAS and PLA are mixed fast muscles with ∼20% oxidative glycolytic type IIA fibers, while the TA and EDL are ∼95% fast glycolytic type IIX or IIB fibers (*44*). Though fiber-type effects were different across muscle examined, this is frequently observed when multiple muscles are analyzed (*43*, *45*, *46*). Specifically, shifts in fiber type composition often are observed under conditions that drive atrophy (e.g. unloading, denervation), which reflects a combination of loading and stress stimuli that differs between conditions (*47*). *Pptc7* KO shifted fibers in the SOL toward slow oxidative (type IIA -> type I) and fibers in the mixed PLA and GAS toward fast glycolytic (type IIA -> type IIB), and these differential responses are likely complex and governed by both loading and metabolic factors. It is also worth noting that non-autonomous factors may influence fiber or tissue properties, as our inducible *Pptc7* KO model promotes gene excision across multiple cell types and tissues, which may enable non-muscle effectors to contribute to the phenotypes reported.

Despite this, the phenotypical deviations seen across *Pptc7* KO muscles do not seem to stem from technical artifacts such as mosaicism, as our genotyping and immunoblotting data suggest robust excision and loss of PPTC7 across all tested muscle types. Instead, several orthogonal pieces of data suggest that dysregulation of BNIP3-mediated mitophagy contributes to the muscle dysfunction seen adult, inducible *Pptc7* KO knockout mice. First, all tested *Pptc7* KO skeletal muscles showed increased BNIP3 protein levels, though with some variability in the amount or the distribution between its monomeric and dimeric forms. Second, the SOL and EDL–two muscles with significant upregulation of at least one BNIP3 species–were profiled by transmission electron microscopy which revealed significantly elevated numbers of autophagosome- or mitophagosome-like structures in *Pptc7* KO tissues, consistent with the model that loss of *Pptc7* elevates mitophagic flux through the upregulation of BNIP3. Recent work suggests that upregulation of BNIP3- and NIX-mediated mitophagy decreases mitochondrial mass but that organelles retain function (*48*), suggesting that, if mitophagy were driving tissue dysfunction, mitochondria may decrease in abundance but remain competent in respiration. Indeed, we obtained significantly lower yields of mitochondria from *Pptc7* KO GAS tissue than from floxed controls, that, when normalized for content, displayed respiratory rates indistinguishable from organelles derived from floxed controls. These data suggest that, in the GAS, there does not appear to be intrinsic organellar dysfunction caused by the loss of *Pptc7*, at least at the level of respiratory capacity. Interestingly, however, *Pptc7* KO mouse embryonic fibroblasts have diminished oxygen consumption that is not fully rescued by simultaneous dual knockout of *Bnip3* and *Bnip3l* (i.e., NIX) (*7*). These data suggest that regulation of BNIP3-mediated mitophagy may be particularly important in skeletal muscle, but that other functions of PPTC7, such as its phosphatase activity, may contribute to the biology of other cell types. Further investigation of the contributions of this multifunctional phosphatase across tissues, developmental states, and organismal contexts will be critical to understand its role in enabling mammalian metabolic function.

Beyond underscoring the complexities of responsiveness to loss of PPTC7 across developmental stages, our data suggest that proper regulation of mitophagy is critical in skeletal muscle, and that responses to its disruption are complex. Indeed, accumulating evidence suggests that mitophagic flux is dynamic and variable across skeletal muscles in physiological and pathological contexts. For instance, studies in mice expressing the mitophagy reporters mt-Keima (*49*) and mito-QC (*50*) have found that muscles with more oxidative capacity, such as the SOL and R-GAS, have low ratios of mitolysosomes relative to total mitochondrial content as compared to the more glycolytic W-GAS or EDL muscles (*51*, *52*), suggesting that glycolytic muscles have higher mitophagic flux than oxidative muscles. Given the known differences in mitochondrial ultrastructure, function, and proteome content across skeletal muscles and fiber types (*53–55*), these data suggest that mitophagy may contribute to or even enable such diversity in organellar composition and content. Furthermore, mouse models and human patients with mitochondrial myopathies display dysfunctional mitophagy which, intriguingly, occurs in a mosaic manner, differing even between adjacent muscle fibers (*56*), suggesting that heterogeneity in skeletal muscle mitophagy may contribute to (or at least persist during) certain pathologies. Consistently, the literature on BNIP3 itself is often contradictory, with some studies demonstrating that elevated levels of BNIP3 lead to muscle dysfunction, including atrophy during cachexia (*30*), fiber atrophy during late-stage genetic disorders such as Pompe disease (*28*), or correlate with worse outcomes during immobilization-induced atrophy (*29*). Other studies, however, suggest that ablation of *Bnip3* does not rescue skeletal muscle function in models of cancer cachexia (*57*), and that its downregulation in the muscles of aged mice can exacerbate atrophy (*58*). Such contradictions likely highlight the complexities of BNIP3 function and/or mitophagy across skeletal muscle types, developmental stages, and disease states.

Given the growing appreciation of variability of mitophagic regulation in skeletal muscle in physiology and pathology (*59*), we propose that careful selection of skeletal muscles to be analyzed, as well as other parameters such as mouse sex, age, and developmental stage, will be critical experimental determinants to normalize in future studies. This importance is highlighted in our study, as different muscles display distinct phenotypes in response to the same genetic perturbation, even within the same mouse. Beyond emphasizing the importance of experimental considerations, our study also suggests that caution should be exercised in augmenting mitophagy as a therapeutic strategy. Though multiple groups have identified mitophagy as protective in skeletal muscle pathologies (*60–63*), our data suggests this process requires fine-tuned regulation across muscle and/or fiber types and developmental stages. Though complex, these data suggest that cells and tissues employ diverse strategies to sense and respond to dysregulated mitophagy and elevated mitochondrial clearance. As such, continued studies of such systems should provide rich new insights into the differential regulation and responsiveness of dynamic mitochondrial behavior in vivo.

## Acknowledgements

We thank Dr. Gerald Dorn III for sharing the global *Bnip3* KO mouse strain (*64*), and the Mouse Genetics Core at Washington University School of Medicine in assisting us with its rederivation. We gratefully acknowledge Gregory Strout and John Wulf II for their assistance in electron microscopy studies conducted at the Washington University Center for Cellular Imaging (WUCCI). We thank Dr. Sangeeta Adak as well as the Animal Model Research Core, the Diabetes Models Phenotyping Core, and the Washington University Musculoskeletal Research Center at Washington University School of Medicine in St. Louis. We thank members of the Niemi laboratory, Dr. Brian Finck (WUSM) and Dr. Jonathan Friedman (UTSW) for critical reading and feedback on the manuscript, as well as for continued discussions and advising on the work.

## Supplementary Materials

Figs. S1 to S5 are featured at the end of the manuscript.

## Author contributions

Conceptualization (T.M.L., N.M.N.), Data acquisition and curation (T.M.L., T.N.M., K.C., S.V., M.M.M., M.F., N.M.N), Formal Analysis (T.M.L., T.N.M., N.M.N.), Funding acquisition (N.M.N.), Methodology (T.M.L., T.N.M., K.C., J.L.A.F., M.F., L.P.S., G.A.M., G.J.P), Resources (K.C., J.L.A.F., L.P.S., G.A.M., G.J.P.), Visualization (T.M.L., T.N.M., N.M.N.), Writing – original draft (T.M.L., G.A.M., N.M.N.), Writing, review and editing (all authors).

## Data, code, and materials availability

All data are available in the manuscript or the supplementary materials. Code/macros for skeletal muscle fiber typing analysis available upon request.

## Methods

### Mouse strains, husbandry, and ethical approval

The mouse strains used in this study are commercially available and/or have been described previously: inducible *Pptc7* KO animals (UBC-Cre^ERT2^;Pptc7^flox/flox^, originally described in (*7*)), global *Pptc7* KO animals (only the E2 mutants were used in this study, originally described in (*6*)); and global Bnip3 KO animals (a kind gift from Gerald Dorn III, originally described in (*64*)).

Tamoxifen-induced knockout was accomplished in the inducible *Pptc7* KO animals in a protocol slightly modified from previously described. Briefly, upon reaching 8 weeks of age, mice were fed tamoxifen-containing chow for ten total days, split across five days with tamoxifen chow and two days with normal chow for two consecutive weeks. Tamoxifen chow was purchase from Envigo (Teklad custom diet TD.130859), which contains 400 mg of tamoxifen citrate per kilogram of diet. To promote feeding, powdered tamoxifen chow was mixed at a ratio of 3:1 tamoxifen chow with ground standard chow (Formulab Diet 5008). 20 grams of mixed diet (i.e., 15 g. Tamoxifen chow and 5 g. Standard chow) were placed into weigh boats, with water added to create a mash. This mash was made freshly and administered to each cage for each of the 10 days of the tamoxifen treatment protocol. On the 6th day of each week, the tamoxifen chow was removed, and standard chow was given to mice for a total of two days. This routine (5 days tamoxifen, 2 days standard chow) was repeated for a total of two consecutive weeks. After tamoxifen administration was complete, mice were fed standard chow for the duration of the experiment. All mice were housed on a 12-hr. light:dark cycle, and were group housed by strain and sex under temperature- and humidity-controlled conditions. Unless noted, all animals received ad libitum access to water and food. All animal work was performed in accordance with ethical regulations for animal research via IACUC approval through Washington University School of Medicine in St. Louis (animal welfare assurance D1600245).

### Genotyping analysis

Each mouse in our study was validated by genotyping analysis. Routine genotyping occurred via one of two methods. In the first, we performed in house genotyping. Tail tips were isolated from each mouse at approximately P14 and genomic DNA (gDNA) was extracted by adapted protocols ((*6*, *7*)). Briefly, the tail tips were placed in microcentrifuge tubes with the addition of 600 μL of genomic lysis solution (20 mM Tris, pH 8.0, 150 mM NaCl, 100 mM EDTA, 1% SDS) supplemented with 5 μL of proteinase K (20 mg/mL) and incubated at 55*C overnight. After incubation and ensuring full digestion of the tail tips, 200 μL of Puregene Protein Precipitation Solution (catalog# 158123, QIAGEN, Hilden, Germany) was added and samples were incubated on ice for 5 minutes. Samples were vortexed for approximately 15 seconds and then centrifuged (16,000 x g) for 6 minutes to pellet the precipitated proteins. The supernatant was removed and placed in a fresh 1.5 mL microcentrifuge tube and supplemented with 600 μL of 100% isopropanol. Sample tubes were inverted approximately 25x and centrifuged (16,000 x g) for 3 minutes to precipitate the gDNA. The supernatant was aspirated and the pellet was washed with 600 μL of 70% ethanol and centrifuged a final time (16,000 x g) for 3 minutes. After aspirating the supernatant again, the pelleted gDNA was allowed to dry overnight at room temperature. After adequate drying, 50-100 μL of ultrapure water (depending on the size of the pellet) was added to each sample, gently mixed, and gDNA was allowed to rehydrate for at least one hour. gDNA was quantified using a Take3 microvolume plate (catalog# TAKE3-SN, BioTek Instruments, Winooski, VT, USA) and BioTek Gen5 software in an Epoch2 microplate spectrophotometer and approximately 100-200 ng of gDNA was added to each genotyping reaction.

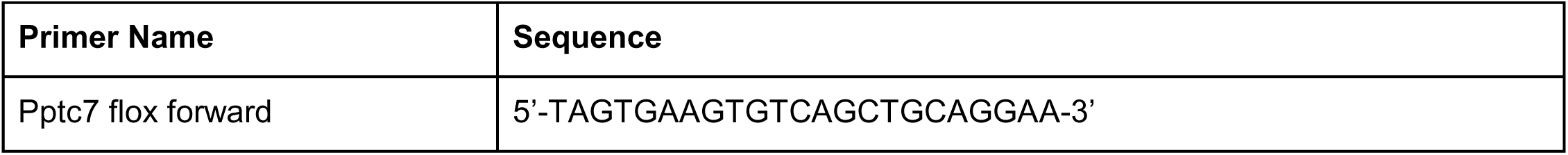
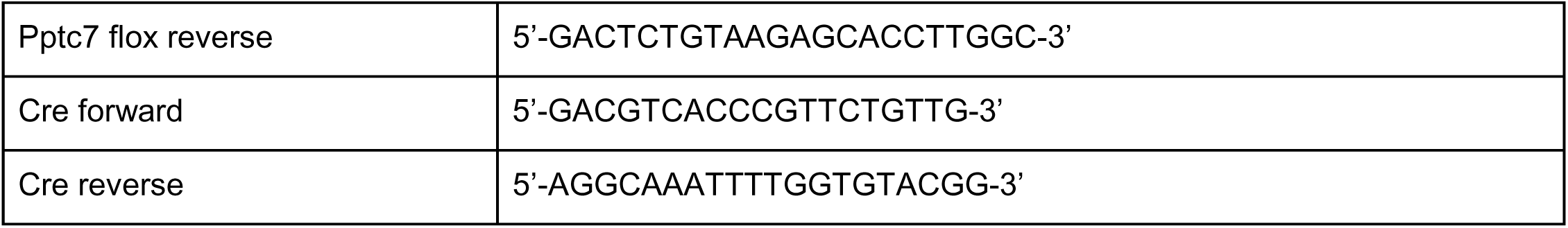

In the second method of routine genotyping, animals were genotyped through commercially available and optimized methods through Transnetyx. In this method, tails were isolated as described above and sent to Transnetyx, where they were processed and genotyped via appropriate strain-specific protocols.

Finally, we performed in house genotyping of tissues derived from select animals to confirm their genotypes and the degree of potential mosaicism across tissues. gDNA from mouse tissues was extracted using the DNeasy Blood and Tissue Kit (catalog# 69504, QIAGEN, Hilden, Germany) using the manufacturer’s instructions. Briefly, tissues were minced into small pieces using scissors in a 1.5 mL microcentrifuge tube. 180 μL of Buffer ATL and 20 μL of Proteinase K (20 mg/mL) were added and the tubes were mixed by vortexing and incubated on a heat block at 56*C overnight, or until the tissues were completely lysed. 200 μL of Buffer AL was added to the samples, vortexed, followed by the addition of 200 μL of 100% ethanol and mixed again by vortexing. The vortexed samples were then transferred into the DNeasy mini spin columns placed into 2 mL collection tubes and centrifuged at 6000 x g for 1 minute at room temperature. The flow-through was discarded and columns were washed with 500 μL of Buffer AW1 and centrifuged at 6000 x g for 1 minute at room temperature. The flow-through was again discarded and columns were washed with 500 μL of Buffer AW2 and centrifuged at 20,000 x g for 3 minutes at room temperature to dry out the filter membranes. The flow-through and collection tubes were discarded and the spin columns were transferred to new 1.5 mL microcentrifuge tubes. 100 μL of Buffer AE was added directly to each membrane and allowed to incubate on the membranes for at least one minute. Each tube was centrifuged at 6000 x g for 1 minute at room temperature to elute the genomic DNA. The gDNA was quantified using a Take3 microvolume plate (catalog# TAKE3-SN, BioTek Instruments, Winooski, VT, USA) and BioTek Gen5 software in an Epoch2 microplate spectrophotometer and approximately 100-200 ng of gDNA was added to each genotyping reaction.

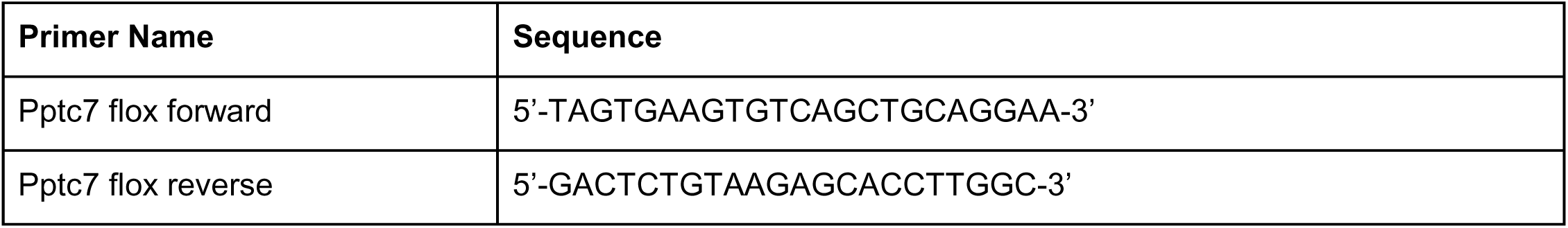

The floxed allele amplifies at a size of 667 base pairs, while the knockout allele amplifies at a size of 164 base pairs. Mosaicism was noted when both alleles amplified from the same gDNA sample.

### Murine body composition analysis and indirect calorimetry

Mice were evaluated for whole body metabolic parameters using indirect calorimetry using a PhenoMaster metabolic cage system (TSE Systems). Mice were single housed for 5-7 days before measurements were taken to acclimate them to this potential stressor. Data were collected across eight cages for a 72-hour period with food accessible ad libitum; the first ∼24 hours of measurements were not used, as animals were acclimating to the metabolic cages. Animals housed at room temperature, which was measured and ranged between ∼70-73 degrees Fahrenheit (i.e., below thermoneutrality for mice). Water and food were given ad libitum with consumption measured through high precision weight sensors within the Phenomaster unit. Data were retrieved from the system and experimental runs for female or male mice were combined using the CalR app. Each combined analysis was then analyzed using the CalR2 app (*65*), with data analyzed and reported as suggested in (*21*). Fat and lean mass were measured using an EchoMRI (EchoMRI LLC) body composition analyzer. At least two measurements for fat and lean mass were taken for each animal, with the average of these measurements reported for each animal.

### Serum and plasma isolation, storage, and analysis

#### Serum

Pleural blood was collected (using a 1 mL syringe with 23G needle) from each mouse following cardiac puncture and centrifuged at 21,100 x g for 10 minutes at 4*C. The supernatant (i.e., the serum) was transferred to a new collection tube and immediately snap-frozen in liquid nitrogen. Serum was stored at -80*C until use.

#### Plasma

1.5 mL microcentrifuge tubes were prepared with 30 μL 0.5 M EDTA along with individual 1 mL syringes with 23G needles flushed with 0.5 M EDTA just before plasma collection. Mice were euthanized according to approved procedures, and pleural blood was collected following cardiac puncture. Microcentrifuge tubes containing blood and EDTA were centrifuged at 1,000 x g for 15 minutes at 4*C. The supernatant (i.e., the plasma) was transferred to a new collection tube and immediately snap-frozen in liquid nitrogen. Plasma was stored at -80*C until metabolomics analysis.

#### Mouse plasma metabolomics

Ultra-high performance liquid chromatography coupled with mass spectrometry (UHPLC/MS) analysis was performed using a Thermo Scientific Vanquish Flex UHPLC system interfaced with a Thermo Scientific Orbitrap ID-X mass spectrometer. Polar metabolites were separated on a HILICON iHILIC-(P) Classic HILIC column (100 × 2.1 mm, 5 µm) equipped with a HILICON iHILIC-(P) Classic guard column (20 × 2.1 mm, 5 µm). The mobile phases consisted of solvent A (20 mM ammonium bicarbonate, 5 µM medronic acid, 0.1% ammonium hydroxide in 95:5 water:acetonitrile) and solvent B (95:5 acetonitrile:water). The column temperature was maintained at 45°C, and metabolites were eluted using a linear gradient at a flow rate of 250 µL/min with the following gradient profile: 0-1 min, 90% B; 12 min, 35% B; 12.5-14.5 min, 25% B; 15 min, back to 90% B. The injection volume was 4 μL for all polar experiments. Data was acquired in positive and negative ion mode with the following settings: spray voltage, 3.5 kV (positive) and −2.8 kV (negative); sheath gas, 50; auxiliary gas, 10; sweep gas, 1; ITT temperature, 300°C; vaporizer temperature, 200°C; mass range, 67–1,000 Da; resolution, 120,000. LC/MS data were processed and analyzed using Compound Discoverer and Skyline software (*66*). Natural-abundance correction of ^13^C was performed with AccuCor (*67*).

#### Measurements of FGF-21 and GDF-15

Serum FGF-21 and GDF-15 levels were assayed using the Mouse/Rat FGF-21 Quantikine ELISA Kit (catalog# MF2100, R&D Systems, Minneapolis, MN, USA) and the Mouse/Rat GDF-15 Quantikine ELISA Kit (catalog# MGD150, R&D Systems, Minneapolis, MN, USA), respectively, according to the manufacturer’s instructions. Briefly, serum samples were thawed completely on ice. 50 μL (diluted 2-fold for FGF-21, non-diluted for GDF-15) of serum was assayed from each mouse and FGF-21 and GDF-15 levels were quantified relative to the standard curves generated from each kit. The positive control solution from each kit was verified to have a concentration of FGF-21 or GDF-15 that fell within the manufacturer’s estimated range.

#### Immunoblotting, tissue processing, and antibodies

Tissues were placed in individual 2.0 mL microcentrifuge tubes with a rounded base (catalog# 1620-2700, USA Scientific, Ocala, FL, USA) along with a 5 mm stainless steel bead (catalog# 01-804-413, Fisher Scientific, Fair Lawn, NJ, USA). The tissues were homogenized in lysis buffer A (50 mM Tris, pH 7.4, 40 mM NaCl, 1 mM EDTA, 0.5% triton-X 100) supplemented with 1X protease inhibitor cocktail (0.5 μg/ml pepstatin A, chymostatin, antipain, leupeptin, and aprotinin) and 1X phosphatase inhibitor cocktail (0.5 mM imidazole, 0.25 mM sodium fluoride, 0.3 mM sodium molybdate, 0.25 mM sodium orthovanadate, and 1 mM sodium tartrate) using the TissueLyser II (QIAGEN, Hilden, Germany) set to 30 Hz and 90 seconds. Tissue lysates were clarified by centrifugation (21,100 x g) for 10 minutes at 4*C and the concentration of each clarified lysate was quantified with the bicinchoninic acid assay kit (catalog# PI23225, Thermo Fisher Scientific, Waltham, MA, USA). Lysates were then snap frozen with liquid nitrogen and stored at -80*C until use. For immunoblotting, lysates (40 μg, unless otherwise specified) were supplemented with 0.2% SDS, 1.0% Triton X-100, and 5X sample buffer (312 mM Tris-Base, 25% wt/vol sucrose, 5% wt/vol SDS, 0.05% wt/vol bromophenol blue, 5% vol/vol β-mercaptoethanol, pH 6.8) and boiled at 95*C for 10 minutes. Following protein denaturation, lysates were run on SDS-PAGE gels with Precision All-Blue Protein Standards (catalog# 1610373, Bio-Rad Laboratories, Hercules, CA, USA) prior to being transferred onto nitrocellulose membranes. Following transfer, all membranes were incubated in a blocking buffer (2% bovine serum albumin) for 30 minutes prior to incubating in primary antibodies (diluted in 2% bovine serum albumin) for periods indicated below. The primary antibodies used in immunoblotting include: anti-PPTC7 (catalog# NBP190654, dilution 1:1,000, 48 h incubation at 4°C; Novus), rodent anti-BNIP3 (catalog# 3769, dilution 1:1,000, 48 h incubation at 4°C; Cell Signaling Technology [CST]), anti-NIX (catalog# 12396, dilution 1:1,000, 48 h incubation at 4°C; CST), anti-GAPDH (catalog# 2118, dilution 1:1,000, overnight incubation at 4°C; CST), anti-GAPDH (catalog# 5174, dilution 1:1,000, overnight incubation at 4°C; CST). After primary incubation, the membranes were washed three times with 1X TBS-T prior to incubation in the fluorophore-conjugated anti-800 rabbit antibody (LiCOR Biosciences, Lincoln, NE, USA), diluted 1:10,000 in 1X TBS-T, for 30 minutes at room temperature. Membranes were washed three times with 1X TBS-T and imaged using a LiCOR OdysseyFC instrument equipped with Image Studio software version 5.2 (LiCOR Biosciences, Lincoln, NE, USA).

#### Skeletal muscle fiber typing and analysis

The gastrocnemius, soleus, and plantaris (also known as the plantarflexor muscles) of the mouse were necropsied fresh and affixed together to cork using tragacanth gum and flash frozen in isopentane cooled by liquid nitrogen. Samples were stored in -80°C until sectioning. At the time of sectioning, frozen muscle bundles were sectioned at the mid-belly axially at 10 μm with a cryostat (Leica Biosystems, Wetzlar, Germany). Sections were allowed to dry on the slides for 1 hour. Sections were then rehydrated using 1X phosphate buffered saline for 10 minutes. Then, sections were immunostained for 1 hour at room temperature protected from light against myosin heavy chain isoforms type I, type IIA, and type IIB (each 1:30 in 2% bovine serum albumin; BA-F8, SC-71, BF-F3; Developmental Studies Hybridoma Bank, Iowa City, IA, USA) and counterstained against laminin (1:400 in 2% bovine serum albumin; ab11575; Abcam, Cambridge, UK). After washing three times with 1X phosphate buffered saline, sections were stained with compatible secondary antibodies (diluted 1:400 in 2% bovine serum albumin; catalog# A21242 for type I, A21120 for type IIA, A21042 for type IIB, and A11011 for laminin; Fisher Scientific, Fair Lawn, NJ, USA) and incubated for 30 minutes at room temperature protected from light. After washing three times with 1X phosphate buffered saline, 100% methanol was added to each slide and incubated for 5 minutes at room temperature protected from light. After washing three times with 1X phosphate buffered saline, sections were covered with a small amount of Fluoromount G (catalog# 50-187-88, Fisher Scientific, Fair Lawn, NJ, USA) and sealed with a coverslip and clear nail polish.

Imaging was performed using a Zeiss Axio Scan.Z1 microscope with a Plan-Apochromat 20X objective equipped with a Hamamatsu Orca Flash digital camera and Zen 3.1 (blue edition) software. To detect the fluorescent antibodies, four channels were actively utilized during imaging: Alexa 405 (to detect type IIA fibers), Alexa 488 (to detect type IIB fibers), Alexa 568 (to detect laminin), and Alexa 647 (to detect type I fibers). Unstained fibers were classified by type IIX. Two distinct sub-images of consistent size with minimal-to-no tissue tearing or folding were chosen for each muscle (e.g., SOL, PLA, R-GAS, or W-GAS) and biological replicate equating to at least 75 total fibers per set of sub-images. Fiber size by fiber type (i.e., cross-sectional area, CSA) was determined using a semi-automated ImageJ macro described previously (*68*). Briefly, the laminin (i.e., Alexa 568) channel is opened for a particular sub-image and adjusted using filters (e.g., Gaussian Blur, Unsharp Mask) and auto-thresholding (e.g., Huang dark) to generate a segmented sub-image with regions of interest (ROIs, i.e., individual fibers) that could be analyzed (size = 200-20000 μm^2^) by the subsequent opening of the fiber type-specific channels. The output ‘Set Measurements’ were ‘Area’ and ‘Fit ellipse’. Individual fibers that had a major divided by minor (i.e., major/minor) value greater than 2.25 were omitted from the CSA analysis as these fibers were considered to be cut off-angle. To ensure no biological replicate was oversampled in comparison to the others, fiber sizes were then ranked from lowest major/minor value to highest major/minor value and the first ‘X’ number of fibers were recorded for the fiber size-by-fiber type analysis. In the soleus, both type I and type IIA fibers included 50 fibers per mouse. In the plantaris, type IIA, type IIB, and type IIX included between 20-50 fibers per mouse. In the red gastrocnemius, type I, type IIA, and type IIB included between 20-50 fibers per mouse, while only 10-35 type IIX fibers could be quantified per mouse. In the white gastrocnemius, type IIB fibers included 50 fibers per mouse. For the fiber distribution analysis, the total number of fibers from each fiber type per set of sub-images were calculated as a percentage.

#### Transmission Electron Microscopy

Animals were transcardially perfused with Mammalian Ringer Solution (catalog# 50-980-245, Fisher Scientific, Fair Lawn, NJ, USA). Soleus and extensor digitorum longus muscles were excised and approximately 1 mm^3^ pieces were immersion fixed overnight at 4*C in a solution containing 2% paraformaldehyde and 2.5% glutaraldehyde in 0.15 M cacodylate buffer with 2 mM CaCl2, pH 7.4. Samples were then rinsed in cacodylate buffer 3 times for 10 minutes each and secondary fixed for one hour in 2% osmium tetroxide in cacodylate buffer with 2 mM CaCl2. Following this, samples were rinsed in ultrapure water 3 times for 10 minutes each and stained overnight in an aqueous solution of 2% uranyl acetate at 4*C. After staining was complete, samples were washed in ultrapure water 3 times for 10 minutes each, dehydrated in a graded ethanol series (50%, 70%, 90%, 100% x4) for 10 minutes in each step, then infiltrated and embedded via a graded series of Epon 812 Resin (Electron Microscopy Sciences, Hatfield, PA) with vacuum and microwave assistance (Pelco BioWave Pro, Redding, CA) into Epon 812 resin. Samples were cured in an oven at 60*C for 72 hours. before ROIs were selected, sectioned at 70 nm, and stained with 2% uranyl acetate (10 minutes), and Reynolds lead (2 minutes).

Imaging was performed using a JEOL JEM-1400 Plus Transmission Electron Microscope operated at 120kV with an AMT (Woburn, MA) Nanosprint15-MkII sCMOS camera. Micrographs were captured at magnifications of 3000x (to view mitochondrial network across muscle fibers) or 10000x (to view individual mitochondrion ultrastructure).

#### High-resolution respirometry of permeabilized fibers and isolated mitochondria

The soleus (SOL) and extensor digitorum longus (EDL) were collected for mitochondrial respiration. The SOL and EDL were blotted on filter paper, weighed, and immediately placed in an ice-cold biopsy preservation buffer (BIOPS) containing 10 mM Ca-EGTA buffer, 0.1 µM free calcium, 20 mM imidazole, 20 mM taurine, 50 mM K-MES, 0.5 mM DTT, 6.56 mM MgCl2, 5.77 mM ATP, 15 mM phosphocreatine, (pH 7.1). Fiber bundles were separated mechanically with two pairs of very sharp forceps, in a small petri dish on ice. After completing fiber separation, the SOL and EDL muscles were transferred quickly into 2 mL of ice-cold BIOPS, containing 20 µL of saponin stock solution (5 mg/mL; final concentration 50 µg/mL) and kept on gentle agitation at 4*C for 30 minutes to achieve permeabilization. Thereafter, permeabilized SOL and EDL muscles were washed twice in 2 mL of BIOPS at 4*C for 15 minutes before proceeding to respirometry.

Mitochondria from the whole gastrocnemius (GAS) muscle were isolated by serial centrifugation. Briefly, the GAS was homogenized in an ice-cold isolation buffer (250 mM sucrose, 10 mM Tris-base, 0.5 mM EDTA, pH 7.4). After centrifuging at 1000 x g for 5 minutes at 4*C to pellet nuclei and cellular debris, the supernatant was collected and centrifuged at 10,000 x g for 10 minutes at 4*C to obtain the crude mitochondrial pellet. The pellet was then washed with isolation buffer and centrifuged at 9,500 x g for 10 minutes at 4*C. Lastly, the resultant crude mitochondrial pellet was recovered in 100 μL of isolation buffer. The protein content was measured using a Pierce 660 nm Protein Assay Kit (catalog# PI22662, Fisher Scientific, Fair Lawn, NJ, USA).

To measure oxygen consumption rates, permeabilized fibers (SOL and EDL) and isolated mitochondria (GAS) were transferred to an Oxygraph-O2K respirometer chamber (Oroboros Instruments, Innsbruck, Austria) where they were suspended in 2 mL of mitochondrial respiration buffer (0.5 mM EGTA, 3 mM MgCl2, 60 mM Lactobionic acid, 20 mM Taurine, 10 mM KH2PO4, 20 mM HEPES, 110 mM sucrose, and 1 g/L fatty acid free bovine serum albumin, pH 7.1) with continuous stirring at 37*C. During experiments, O_2_ concentration was maintained within the range of 250-400 nmol/mL. Mitochondrial respiratory capacity and function were assayed by the addition of saturating concentrations of pyruvate (5 mM), malate (2 mM), ADP (2.5 mM) and succinate (10 mM).

#### Citrate synthase (CS) assays

Citrate synthase activity was assayed as previously described (*4*). Briefly, clarified tissue lysates were normalized to 1 μg/μL using lysis buffer A (50 mM Tris, pH 7.4, 40 mM NaCl, 1 mM EDTA, 0.5% triton-X 100) and kept on ice until assay set-up. Before assay set-up, a fresh 10 mM solution of oxaloacetate (catalog# O4126, Sigma-Aldrich, St. Louis, MO, USA) was made. 5 μg of tissue lysate was added to a 190 μL reaction containing 100 mM Tris, pH 7.4, 100 μM oxaloacetate, 100 μM 5,5 dithio-bis-(2-nitrobenzoic acid) (DTNB, catalog# 22582, Fisher Scientific, Fair Lawn, NJ, USA) in quadriplicate. In three of the four replicates for each sample, reactions were started by the addition of 10 μL of 6 mM acetyl-CoA (catalog# ACOA-RO, Sigma-Aldrich, St. Louis, MO, USA). DTNB absorbance (A412) was monitored with a BioTek Cytation 5 Cell Imaging Multimode Reader and BioTek Gen5 software on an Epoch2 microplate spectrophotometer. Measurements were taken each minute for a total run time of 30 minutes. Relative rates were calculated using the maximum velocity function (on the BioTek Gen5 analysis software) with the background signal subtracted (from the no acetyl-CoA added well) followed by normalization to the average *Pptc7*^flox/flox^ samples.

#### Statistical analysis

Statistical analysis was performed using Prism (GraphPad, version 10.4.2), with specific statistical tests listed within figure legends for associated data. For ANCOVA analysis for metabolic cage work, statistical analysis was performed using the CalR2 app (*65*). For volcano plots associated with metabolomics data, statistical analysis was performed in Microsoft Excel. Graphs were imported into Affinity Designer 2 (Serif, version 2.6.3) for final figure generation.

