## Supplemental Figures for "Sustained loss of *Pptc7* triggers variable skeletal muscle dysfunction and diminished body mass through dysregulation of BNIP3"

**Supplemental Figure 1**

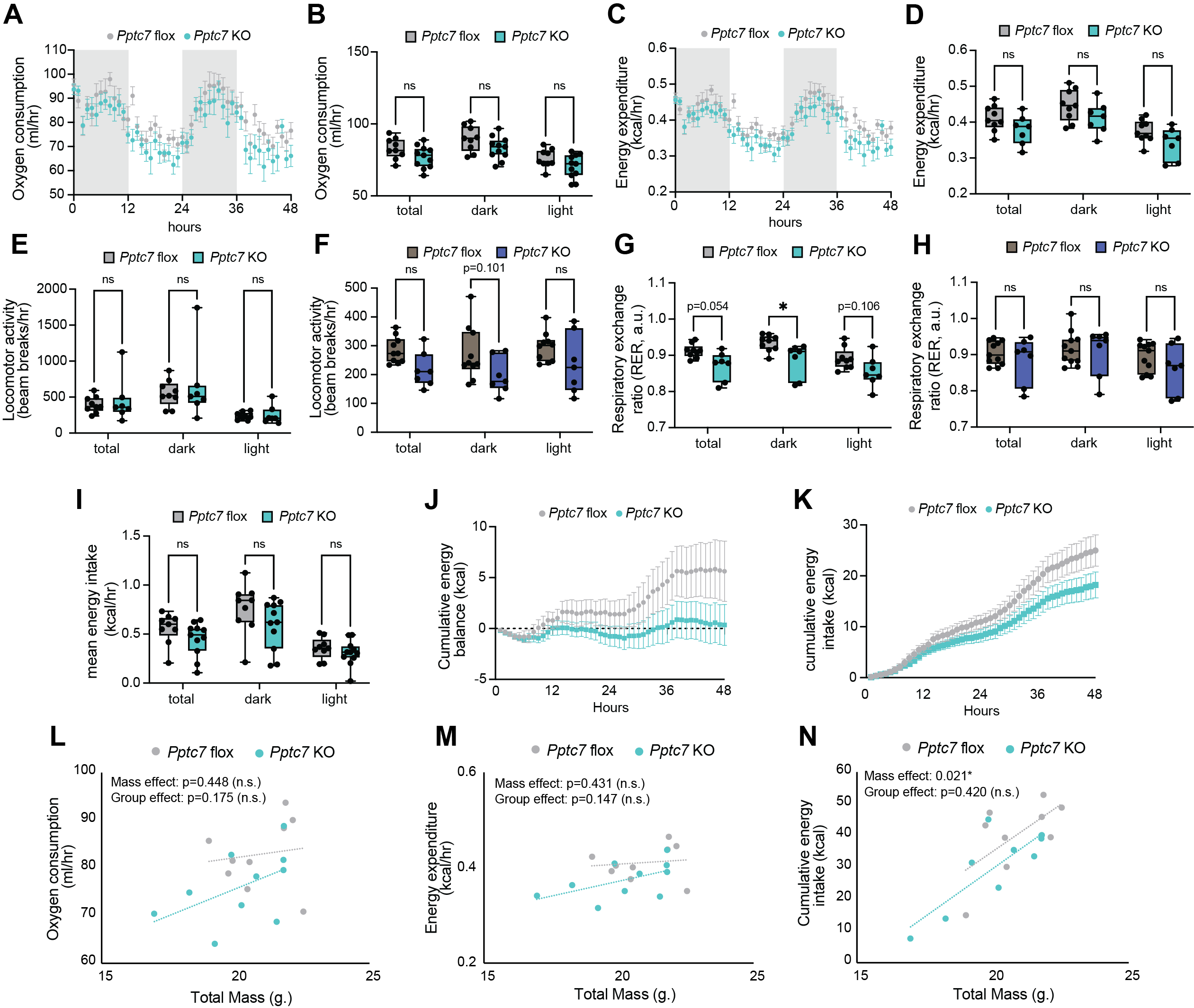

**Supplemental Figure 1**. *Female mice do not demonstrate compromised respiration in response to inducible Pptc7 KO*. **A**., **B**. Oxygen consumption rates for control (*Pptc7* flox) and experimental (*Pptc7* KO) female animals. A. 48-hour average of oxygen consumption rates from female *Pptc7* flox and *Pptc7* KO animals. Error bars represent S.E.M. B. Oxygen consumption rates shown in A. analyzed by circadian parameters for total day averages as well as those during the dark and light phase. n.s. = not significant, Ordinary Two-way ANOVA. **C**., **D**. C. Energy expenditure rates from female *Pptc7* flox and *Pptc7* KO animals. Error bars represent S.E.M. D. Energy expenditure rates shown in C. analyzed by circadian parameters for total day averages as well as those during the dark and light phase. n.s. = not significant, Ordinary Two-way ANOVA. **E., F**. Locomotor activity, as reported in beam breaks/hour, for female (E.) and male (F.) mice. n.s. = not significant, Ordinary Two-way ANOVA. **G., H**. Respiratory exchange ratio for female (G.) and male (H.) mice. * = p<0.05, n.s. = not significant, Ordinary Two-way ANOVA. **I**. Mean hourly energy intake rates for female mice analyzed by circadian parameters for total day averages as well as those during the dark and light phase. n.s. = not significant, Ordinary Two-way ANOVA. **J**. Cumulative energy balance for female *Pptc7* flox and *Pptc7* KO animals over a 48-hour period; error bars represent S.E.M. **K**. Cumulative energy intake for female *Pptc7* flox (n=11) and *Pptc7* KO (n=7) animals over a 48-hour period; error bars represent S.E.M. **L**.-**N**. ANCOVA analysis of weight-dependent parameters such as oxygen consumption rates (L.), energy expenditure rates (M.), and cumulative energy intake (N.) for *Pptc7* flox and *Pptc7* KO female mice. Statistical analysis performed through the CalR2 app, with resulting p-values reported. For boxplots, values between the 25^th^ (bottom) and 75^th^ (top) percentile are displayed, with the median reflected by the line within; whiskers reach the minimum and maximum values.

**Supplemental Figure 2**

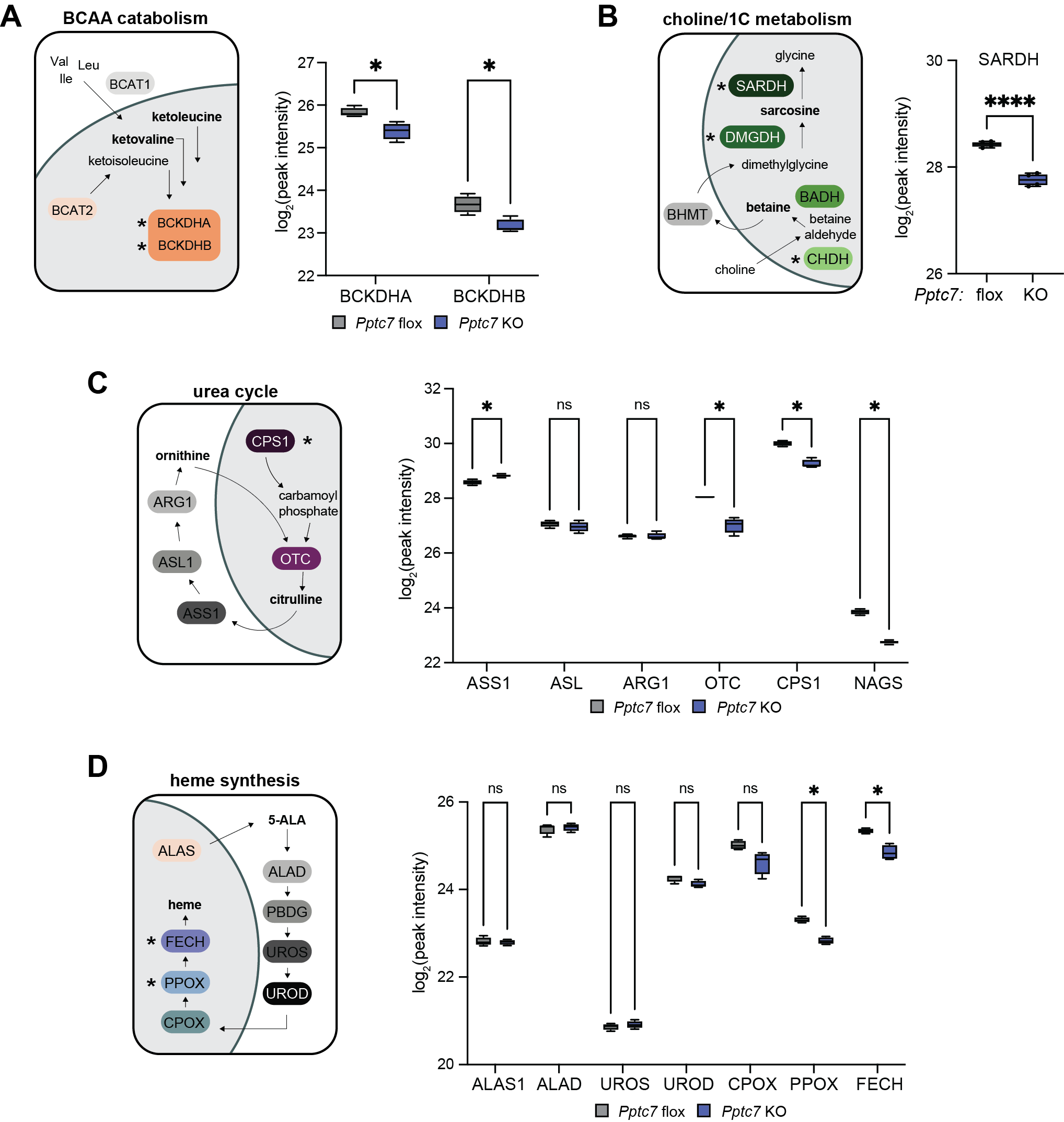

**Supplemental Figure 2**: *Proteomics analysis reveals select decreases in liver proteins that may contribute to alterations in circulating metabolites*. A.-D. Previously collected proteomics data on inducible *Pptc7* KO mouse liver (*7*) reanalyzed for proteins in metabolic pathways that are altered in circulating plasma of this mouse model. A., BCAA proteins, B. SARDH, C. urea cycle proteins, D. heme proteins. Data analyzed with multiple unpaired t tests using the Holm-Šídák method for multiple comparisons. * = p<0.05, n.s. = not significant. Error bars reflect S.D.

**Supplementary Figure 3**

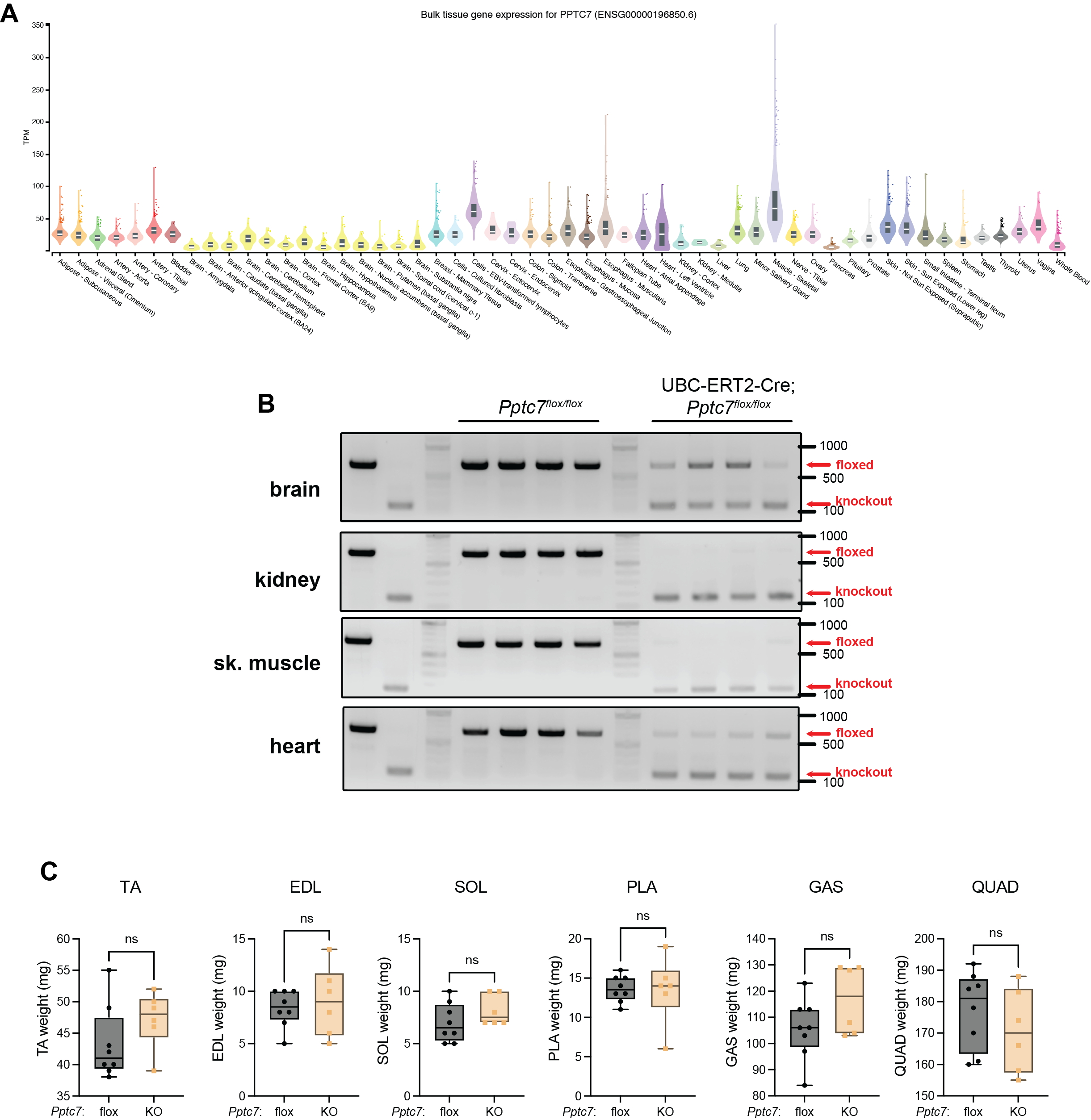

**Supplemental Figure 3**: *Despite high expression and complete excision, Pptc7 knockout females do not display significant skeletal muscle dysfunction at the level of muscle weight*. **A**. GTEx data showing high PPTC7 expression in skeletal muscle (purple) versus other tissues. Data downloaded June 10, 2026. **B**. Genotyping analysis of brain, kidney, skeletal muscle, and heart tissues from *Pptc7* floxed (left) and KO (right) animals. The floxed allele amplifies a larger product than the KO allele, which are both annotated with a red arrow. **C**. Muscles from female *Pptc7* KO animals do not display significant differences in mass relative to their floxed littermate controls. n.s. = not significant, Welch’s t test. For boxplots, values between the 25^th^ (bottom) and 75^th^ (top) percentile are displayed, with the median reflected by the line within; whiskers reach the minimum and maximum values.

**Supplemental Figure 4**

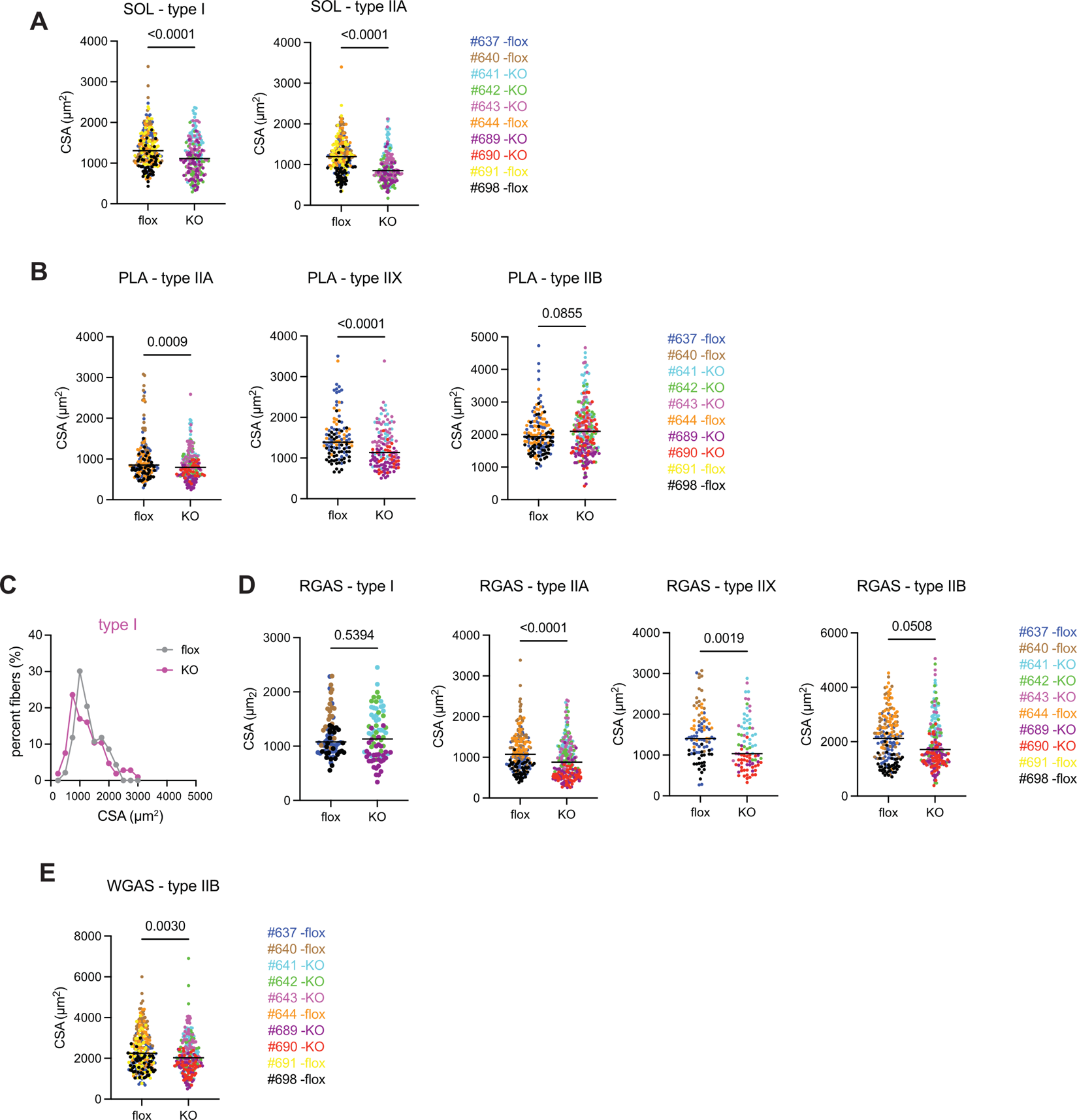

**Supplemental Figure 4**: *Cross sectional areas of various fiber types in Pptc7 KO skeletal muscles*. **A.**, **B**., **D**., **E**. Cross sectional area (CSA) of fiber types represented in the SOL (A.), PLA (B.), RGAS (D.), and WGAS (E.). Fiber type is annotated above graph; fibers not listed for each muscle are not present in sufficient numbers for quantification. On each graph, fibers quantified from each animal are represented as different colored dots; animal number and genotype listed at right. n.s. = not significant, *** = p<0.001, **** = p<0.0001. Welch’s t test. **C**. Histogram distribution of fibers quantified from type I *Pptc7* flox and *Pptc7* KO male mice aged to 20 weeks post-tamoxifen treatment.

**Supplemental Figure 5**

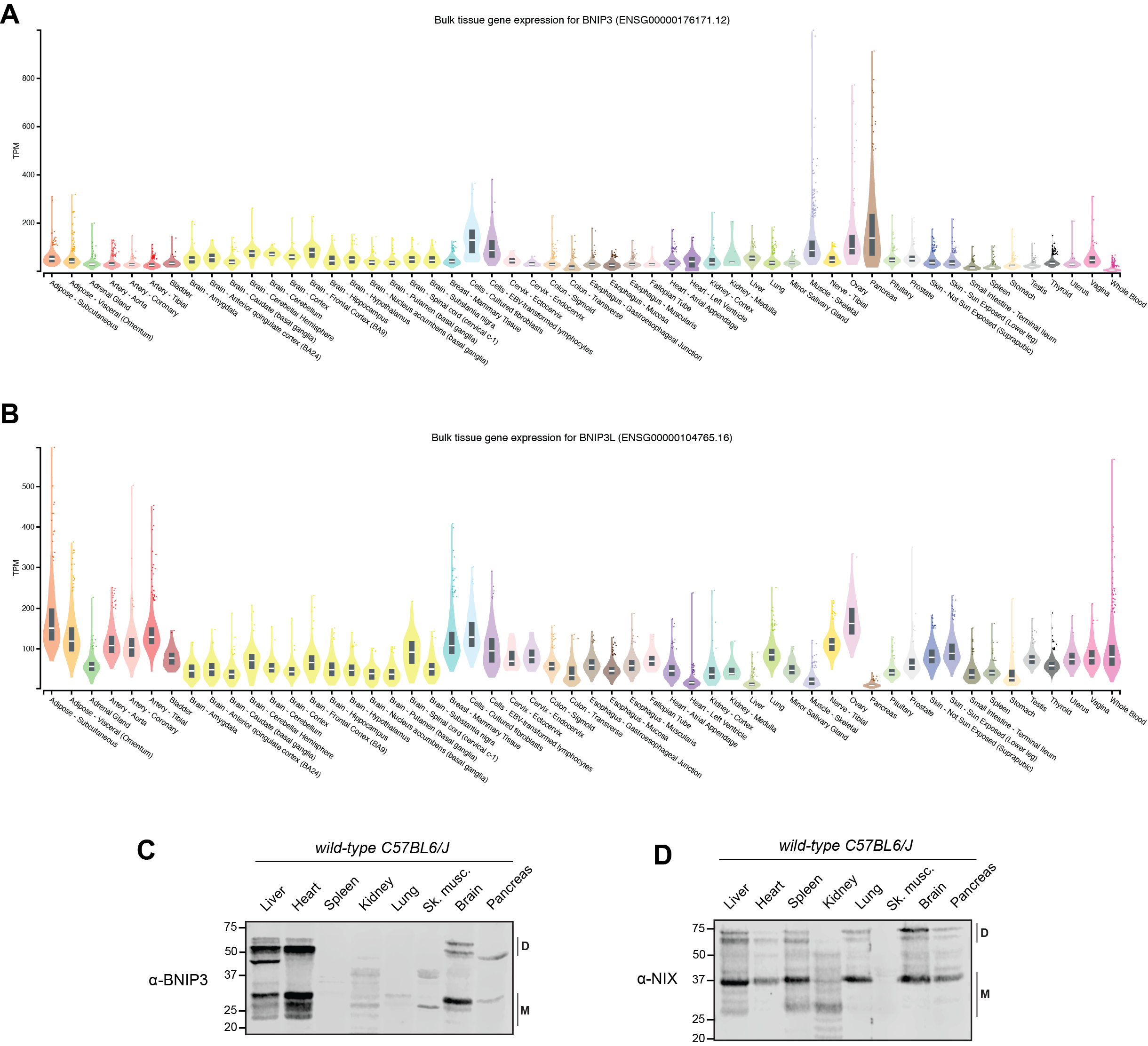

**Supplemental Figure 5**: *Bnip3 expression is higher in skeletal muscle than Bnip3l (i.e., NIX) expression*. **A**., **B**. GTEx data showing high Bnip3 (A.) expression in skeletal muscle (purple) versus other tissues, whereas Bnip3l (i.e., NIX) has relatively low expression in skeletal muscle (purple, B.) Data downloaded June 10, 2026. C., D. Western blot analysis of endogenous BNIP3 (C.) and NIX (D.) expression in a male wild-type C57BL6/J mouse. M = monomeric species, D = dimeric species.
